# Use of IL-6-elafin genetically modified regulatory macrophages as an immunotherapeutic against acute bacterial infection in the lung

**DOI:** 10.1101/2020.07.22.214684

**Authors:** S. Kheir, B. Villeret, I. Garcia-Verdugo, JM Sallenave

## Abstract

**Background:** *Pseudomonas aeruginosa (P.a)* infections are a major public health issue in ventilator-associated pneumoniae, cystic fibrosis and chronic obstructive pulmonary disease (COPD) exacerbations. This bacterium is multidrug resistant and there is an urgent need to develop new therapeutic approaches.

**Objective:** Evaluate the effect of direct pulmonary transplantation of gene-modified (elafin and IL-6) syngeneic macrophages in a mouse model of acute of *P.a* infection.

**Methods:** Wild type (WT) or Elafin-transgenic (eTg) alveolar macrophages (AMs) or bone marrow derived macrophages (BMDMs) were recovered from broncho-alveolar lavage or generated from WT or eTg mice bone marrow. Cells were modified with adenovirus IL-6 (Ad-IL6), characterized *in vitro* (RNA expression, protein secretion, surface markers) and transferred by oropharyngeal instillation in the lungs of naïve mice. The protective effect of the transferred macrophages was assessed during *P.a* acute infection (survival studies, mechanistic studies of the inflammatory response).

**Results:** We show that the transfer in the lung of a single bolus of syngeneic AMs or BMDMs genetically modified with IL6 and Elafin provided protection in our pneumonia *P.a*-induced model. Mechanistically, Elafin-modified AM had an IL-6-IL-10-IL-4R-IL-22-antimicrobial molecular signature which, in synergy with IL-6, conferred, post-transfer, a regulatory phenotype to the alveolar unit.

**Conclusion:** Here we introduce an immunotherapy approach employing gene-modified syngeneic macrophages to target bacterial airway infections. The absence of adverse events during our experiments suggests that our approach is well tolerated and supports the feasibility of translating this therapy to patients suffering from lung acute bacterial infections.

## Introduction

*Pseudomonas aeruginosa (P.a)* is a pathogen causing significant morbidity and mortality, in particular in hospital patients undergoing ventilation and in patients with cystic fibrosis [1][2]. Treatment of lower respiratory tract *P.a* infections becomes particularly problematic since *P.a* is resistant to standard or reserve antibiotic therapy [3]. Indeed, the most recent global priority list of antibiotic-resistant bacteria to guide research, discovery and development of new antibiotics from the WHO, states carbapenem-resistant *P. aeruginosa* as one of the most critical pathogens for which new treatment options are urgently required [4]. Our group has recently shown that *P.a-*derived Elastase B (LasB) degrades the cytokine interleukin (IL)-6 and the antimicrobial molecule Elafin and down regulates a lung epithelial-IL-6-antimicrobial-repair pathway [5], which results in a stronger inflammatory response and a higher mortality rate in a murine model of lung infection. Conversely, we demonstrated that genetic over-expression of a combination of IL-6 and elafin/trappin-2 (a mucosal antimicrobial/anti-elastase molecule) was protective in that same model [5].

Because that approach involved the direct lung instillation of a replication-deficient adenovirus vector (Ad-m-IL-6), by essence more trophic for the epithelium, and because the use of direct instillation of Ad-vectors may be problematic for therapeutic applications, we reasoned that the alveolar macrophage (AM), also targeted by *P.a* LasB [6], may be an ideal vessel for the transfer of this protective activity. Indeed, compared to the direct instillation of viral vectors, a cell therapy approach has the advantage of being immunologically relatively ‘silent’, and has indeed successfully being used in SCID patients, where lymphocytes and stem cells have been efficiently corrected using lentiviruses [7][8]. Furthermore, Ad-vectors offer the advantages, over other vectors such as lentiviruses, to remain episomal in the nucleus, and therefore to not require integration into host chromosomes to deliver their genetic cargo [9].

Crucially, AMs represent the most abundant myeloid cells in alveolar spaces [10][11]. They are critical regulators in the maintenance of immunologic homeostasis [11] in the respiratory tract and play a key role in the initiation and resolution of the immune response in the lungs [12][13]. Given their importance, different studies have focused on macrophage-based immunotherapy approaches that involve direct pulmonary macrophage transplantation. Notably, in a mouse model of hereditary pulmonary alveolar proteinosis (hPAP) characterized by a disruption of the GM-CSF receptor gene *Csf2*, the disease was corrected following the administration of WT macrophages directly in the lungs [14]. Also, a recent study has shown that intra-pulmonary transplantation of bioreactor-derived induced pluripotent stem cells-macrophages (iPSC-Mac), rescue mice from *P.a-*mediated acute infections of the lower respiratory tract [15].

Here, we demonstrate that the transfer in the lung of a single bolus of syngeneic AMs or BMDMs genetically modified with IL6 and Elafin provided protection in our pneumonia *P.a*-induced model. Mechanistically, we demonstrate that prior to transfer, the Elafin-modified AM had an ‘IL-6/IL-10/IL-4R/IL-22/antimicrobial’ molecular signature which, in synergy with IL-6, conferred, post-transfer, a regulatory phenotype (decrease in iNOS, increase in Arg-1, Ym-1, IL-4Ra, IL-10) to the alveolar unit. Importantly, this regulatory activity did not require the presence of innate or adaptive lymphocytes *in vivo*, but involved an AM-epithelial cellular interplay.

## Materials and Methods

### A-*In vivo* experiments

#### Animals

Seven-to ten-week-old male C57BL/6J WT and age-matched Elafin transgenic (eTg) mice were purchased from Janvier labs (Le Genest-Saint-Isle, France) or generated by our group [16] and bred in house, respectively. Rag1, gamma c double KO mice were a gift from Dr J. Di Santo (Institut Pasteur, Paris). Animals were kept in a specific pathogen-free facility under 12h light/dark cycles, with free access to food and water. Procedures were approved by our local ethical committee and by the French Ministry of Education and Research (agreement number 04537.03).

#### Adoptive transfer and PAO1 Infection

Mice were anaesthetized using intraperitoneal injection of 100 µL physiological saline (10% Ketamine, 10% Xylazine). For intratracheal adoptive transfer of AMs or BMDMs, 50 µL of cell suspension were instilled through the oro-pharynx using an air-filled syringe (500 µL), as described previously (5-6). PAO1 infections were performed by intranasal inhalation of 40 µL of bacterial suspension. During lethal infection experiments and survival studies (≥ 10^7 CFU/ mouse), animals body weight was monitored daily. Alternatively, during mechanistic studies and sub lethal infection experiments (≤ 5×10^6 CFU/mouse), animals were euthanized 16 hrs, 3 or 5 days post infection with an overdose of Euthasol ® Vet (Sodium pentobarbital). Tracheas were cannulated, lungs were washed with 1ml of sterile PBS and bronchoalveolar lavage fluids (BALF) were centrifuged (2000 rpm, 10 min). Supernatants were aliquoted and kept at -80°C for ELISA protein dosage and recovered cells were washed and analyzed by flow cytometry (FACS) or cytospin centrifugation followed by May-Grünwald-Giesma (MGG) staining using the Diff-Quick staining kit (Medion Diagnostics).

### B-*Ex vivo* experiments

#### Primary alveolar macrophages

Primary alveolar macrophages (AMs) were isolated from WT and eTg mice by BAL as described above. BALFs were centrifuged (2000 rpm, 10 min, 4°C) and cell pellets were re-suspended in RPMI-Glutamax (10% FCS, 1000 U/ml Penicillin, 100 µg/ml Streptomycin). Cells were cultured for 16 hrs (37°C, 5% CO_2_) in 48 wells Corning ® Costar® culture plates (250,000 cells/well) prior to stimulation or infection.

#### Bone marrow-derived macrophages (BMDM) generation

Bone marrow was extracted from mice femurs and cells were washed with PBS, centrifuged at 1,400 rpm for 7 min (4°C). Pelleted cells were then re-suspended with lysis buffer (Gibco) during 2 min at room temperature to lyse red blood cells. After another wash, cells were seeded in complete DMEM medium (10% FCS, 1000 U/ml Penicillin, 100 µg/ml Streptomycin) containing 30 ng/mL of mouse M-CSF (Peprotech). After two successive medium replacement (at D3 and D6), macrophages were detached, washed and re-suspended in cold sterile PBS at day 9 and kept on ice before further use. For *in vitro* experiments, cells were seeded in 24 well culture plates (500,000 cells/well) in RPMI-Glutamax (10% FCS, 1000 U/ml Penicillin, 100 µg/ml Streptomycin) for 12 hrs prior to stimulation or infection. For *in vivo* adoptive transfer experiments, BMDMs density was adjusted to 10^6 cells/50 µL in sterile PBS before oro-pharyngeal instillation.

#### Total splenocytes proliferation assay

The spleen of a WT C57/Bl6 mouse was harvested, processed and homogenized into a single cell suspension through mechanical disruption. Briefly, the spleen was grinded and filtered on a 40 µM sterile filter. The filter was washed with DMEM-Glutamax (10% FCS, 1000 U/ml Penicillin, 100 µg/ml Streptomycin) and the recovered cells were centrifuged (2000 rpm, 4°). Pelleted cells were re-suspended for 5 minutes at room temperature with 1ml ACK Lysis buffer to lyse red blood cells. Lysis reaction was stopped by adding 30 ml of sterile PBS 1X and cells were centrifuged (2000 rpm, 4°C). Pelleted cells were re-suspended at 2×10^7/ml in warm PBS and incubated for 10 min at 37°C with 2,5 µM Carboxyfluorescein succinamide ester (CFSE). Labelling was quenched with three washes of ice cold DMEM-Glutamax (10% FCS, 1000 U/ml Penicillin, 100 µg/ml Streptomycin). The CFSE-labeled cells were seeded in a 96 wells culture plate (10.10^4 cells / well) in DMEM-Glutamax (10% FCS, 1000 U/ml Penicillin, 100 µg/ml Streptomycin) supplemented with 30 U/ml of mouse rIL-2. 2µL of DynaBeads® Mouse T-activator CD3/CD28 [Gibco Life Technologies] were added in each well except for a control well (negative control). 100 µL of PBS (control) of BALF were then added into the wells and cells were left in culture for 72 hours. Cells were then harvested, washed with FACs buffer (PBS-1% FCS) and CFSE intensity was analyzed by flow cytometry.

### C-*In vitro* experiments

Mouse alveolar epithelial cell line CMT-2 [CMT64/61, European Collection of Authenticated Cell Cultures, ECACC] and mtCC-DJS2, an immortalized adult Clara cell line [17], a kind gift from Dr DeMayo (Baylor College of Medicine, Texas, USA), were cultured in DMEM-Glutamax (10% FCS, 1000 U/ml Penicillin, 100 µg/ml Streptomycin). Cells were placed for 12 hrs. (37°C, 5% CO_2_) in 24 wells Corning ® Costar® culture plates (500,000 cells/well) prior to stimulation.

#### Alveolar macrophages cell line MPI

Alveolar macrophages (cell line MPI [18]) were cultured in RPMI-Glutamax (10% FCS, 1000 U/ml Penicillin, 100 µg/ml Streptomycin), supplemented with 30 ng/ml GM-CSF (Peprotech). Cells were placed for 16 hrs (37°C, 5% CO_2_) in 48 wells Corning ® Costar® culture plates (250,000 cells/well) prior to stimulation or infection.

#### BMDM and CMT-2/DJS-2 epithelial cells co-culture

100,000 of WT or eTg BMDMs pre-infected with Ad-Null or Ad-IL6 were seeded on 12mm diameter Transwell® inserts of 0,4 µM pore size and inserted into 24 wells culture plate, where CMT-2 or DJS-2 had previously been seeded in the lower compartment. Co-cultures were left untreated overnight and then BMDMs were infected with WT PAO1 (MOI 0,5) for 6 hours. Epithelial cell layers were then isolated for RNA analysis.

#### Adenovirus constructs and adenovirus infection

The control Ad-Null [19], Ad-mIL6 [20] and Ad-hElafin [21] constructs were used to transiently overexpress murine IL-6 and human elafin.

Cells (AMs and MPI) were washed 3x with sterile PBS and infected for 6 hrs with the different Ad constructs with a multiplicity of infection (MOI) of 25 in serum free RPMI-Glutamax. The wells were washed, and cells were placed overnight in serum and antibiotics-free fresh RPMI-Glutamax prior to further stimulation or infection with PAO1.

#### *Pseudomonas aeruginosa O1* (PAO1) strain and macrophage infection

PAO1 strain [ATC 15692] was kept in freezing medium (50% Luria Broth LB, 50% glycerol) and stored at -80°C until use. Before infection experiments, PAO1 strain was grown overnight in LB broth in a rotating incubator (200 rpm, 37°C). The bacterial suspension was then diluted in serum and antibiotic free RPMI medium and the optical density (O.D) was measured at 600 nm every two hours until logarithmic growth phase was reached (0,1 < O.D < 0,3) [an O.D of 0,1 is equivalent to a bacterial concentration of 7,7×10^7 CFU/ml]. Bacterial concentration was then adjusted to 1,25×10^6 CFU/ml and cells were infected for 4 hrs with 100 µL of bacterial suspension per well, which corresponds to a MOI of 0.5.

For *in vivo* infection experiments, after the bacterial concentration was determined, the bacterial suspension was centrifuged (5000 rpm, 4°C, 30 minutes) and the pellet re-suspended in sterile PBS at the desired CFU count.

#### Measurement of PAO1 clearance by macrophages (AMs and MPI)

After 4-hour infection, supernatant from each well were recovered and kept on ice. Cells were then lysed with 400 µL of sterile PBS-0,1% Triton X100. Cell lysates were pooled to the supernatant of the corresponding well and serial dilutions were performed. 100 µL of every dilution were spread on LB agar plates and kept overnight at 37°C for CFU count. Bacterial clearance was calculated by normalizing the CFU count at 4h post infection to the CFU count of a control well (without any macrophages) that represents a 0% clearance condition, according to the following formula:

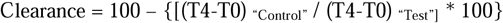

#### RNA extraction, reverse transcription and q-PCR

RNA isolation from cells was performed using PureLink® RNA Mini Kit (12183018A, Ambion, Life technologies), following the manufacturer’s instructions. Briefly, lysates were mixed with 70 % ethanol and loaded onto a silica-membrane column. After different washes, total RNA was eluted in DNase-RNase-free water and stored at -80°C until use after the concentration was quantified using a NanoDrop spectrophotometer.

DNase treatment was performed prior to Reverse transcription polymerase chain reaction (RT-PCR) using RNAse-free DNase I (Roche) at 37°C for 10 min. DNase was then inactivated by increasing the temperature to 70 °C for 10 min. Complementary DNA (cDNA) was synthesized from total RNA (500 ng) using M-MLV Reverse Transcriptase (Promega) as per the supplier’s protocol (1 hr. at 37°C followed by 10 min at 70 °C). Real-time quantitative PCR was performed in a total volume of 15 μL using 2x Fast SYBR® Green Master Mix (Life Technologies), 2 μL of diluted cDNA, 2 μmol forward primer, 2 μmol reverse primer in a 96-well plate. PCR was run with the standard program: 95°C 10 min, 40 times of cycling 95°C 15 sec and 60°C 1 min in a 96-well plate. Results are shown as relative quantity of mRNA copies to the determined control condition with 18s expression used as internal control. These values were calculated according to the comparative ΔΔCt method with:

Relative expression (Fold increase) = 2^ ^(-^ΔΔCt) ; ΔΔCt= ΔCt _Condition -_ ΔCt _Control_ ; ΔCt= Ct _target_ gene – Ct _18s ._ The gene primers used are listed in the Supplemental Material.

### D-Flow cytometry analysis

C57/Bl6 mice lung epithelial cells preparation was as described in Raoust et al (22). For FACS analysis, cells (from lung extracts, BALs, isolated AMs or BMDMs) were first incubated (10 min, 4°C) with a cocktail of a viability dye and Fc Block antibody. Cells were washed (2000 rpm, 5 min) with FACS buffer (PBS-2% FCS), then incubated (30 min, 4°C) with a cocktail of cell surface conjugated antibodies. For intracellular staining, cells were then permeabilized and stained using eBioscience™ Foxp/3 Transcription Factor Staining Buffer Set according to the manufacturer’s protocol. Finally, cells were washed and the pellets were re-suspended in FACS buffer for data acquisition. Data were acquired the same day using LSR Fortessa cytometer (BD Biosciences) with BD FACSDIVA™ software and analyzed with FlowJo (Treestar, OR). The antibodies used for FACS analysis are listed in the Supplemental Material.

### E-Statistical analysis

Data were analyzed with GraphPad Prism Software 6.03. Statistical analysis was performed using either a non-parametric test (Kruskal-Wallis and Dunn’s posttest), or one-way or two-way ANOVA followed by the appropriate multi-comparison post hoc Tukey’s test.

Survival curves in murine model experiments were plotted using Kaplan-Meier curves and statistical tests were performed using the Log-rank (Mantel-Cox) test.

Differences were considered statistically significant when *p* was <0,05 and were labeled as follows: * *p < 0,05; ** p < 0,01; *** p < 0,001; **** p < 0,0001*

## Results

### IL-6 & elafin over-expression modifies cytokines & antimicrobial peptides production and the polarization profile of AMs

We first set up to determine if IL-6 and elafin expression was able to polarize AMs. Unsurprisingly, as previously demonstrated (17), the protein elafin is not expressed in WT C57/Bl6 mice, and was only detected in primary alveolar macrophages (AMs) of elafin transgenic mice (Fig. 1A-B). In un-infected AMs, furthermore, elafin and IL-6 were shown to up-regulate each other : indeed, when compared to controls (WT-AMs), elafin-expressing macrophages (eTg-AM have higher IL-6 levels at steady state (Fig 1 panels C-D). As a mirror image, Ad-mediated IL-6 up-regulated elafin expression in eTg AMs (Fig 1A-B).

**Fig 1.**
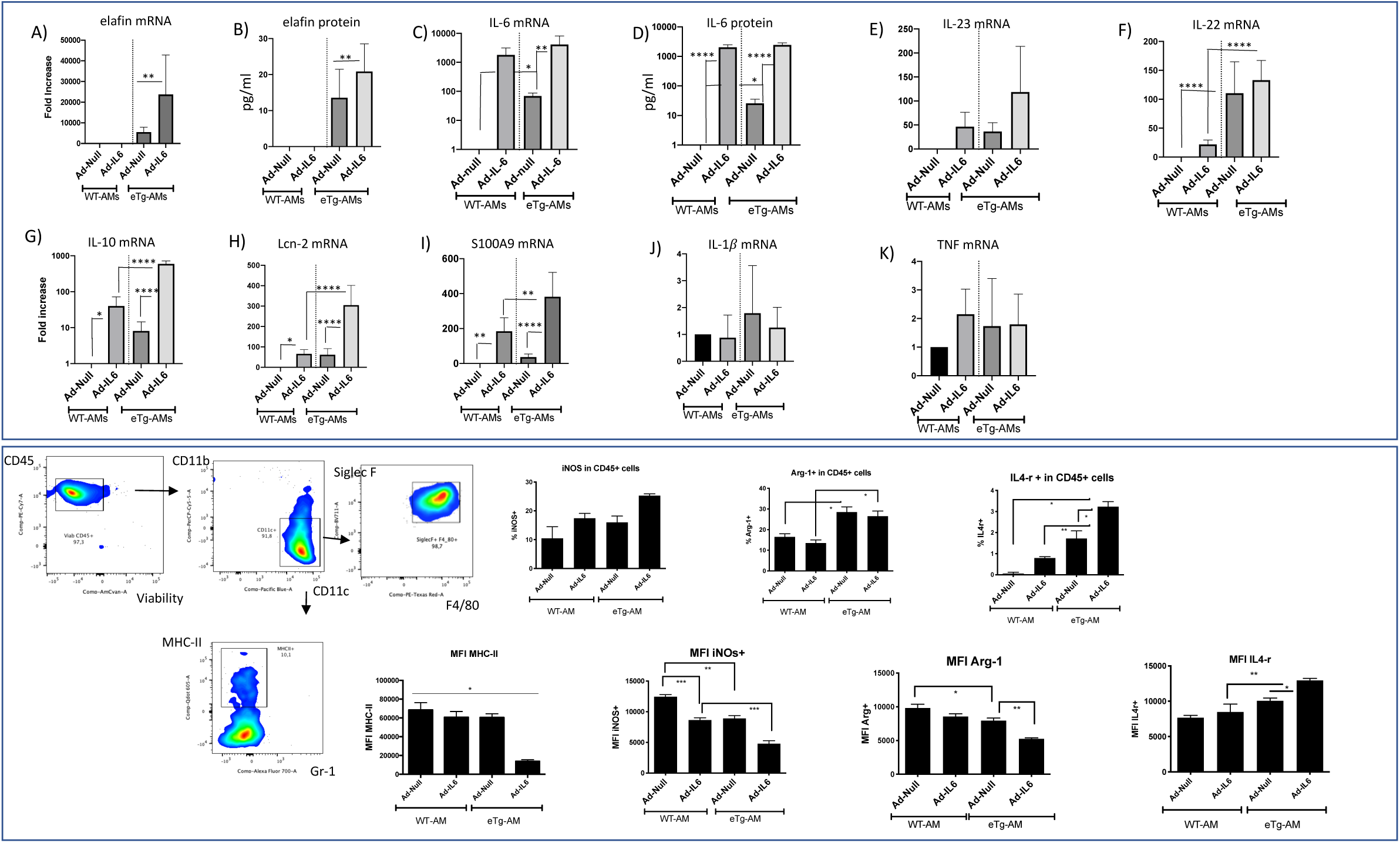
Elafin and IL-6 modify the transcriptional profile and polarization of alveolar macrophages. Upper panel: WT or eTg primary alveolar macrophages (WT-AM or eTg-AM) recovered from mice BALs were transfected for 6 hrs with adenovirus (MOI 25) Ad-Null or Ad-IL-6 and left overnight. Supernatants were then used for IL-6 and elafin protein levels measurements (ELISA). In parallel, cells were then lyzed and the cell lysate were used for mRNA quantification by qPCR according to the ΔΔCt method with Relative expression (Fold increase) = 2^ ^(-ΔΔCt)^ The relative expression, normalized to the housekeeping gene 18s, of elafin, cytokines and antimicrobial peptides mRNA, as well as IL-6 and elafin protein levels in supernatants (measured by ELISAs) are presented. Data show mean +/-SEM (n=4 independent experiments). Statistical significance: ANOVA, multiple comparison, Tukey’s test, * : p < 0.05 ; **: p<0.01; *** :p<0.001; ****: p<0.0001).Lower panel :WT-AM and eTg-AMs were recovered by BALs as above, and were washed with FACS buffer (PBS-FCS 1%) for FACS staining with a viability dye and Fluorochrome-conjugated antibodies against the surface markers CD45, CD11b, CD11c, Gr-1 (Ly6c, Ly6G), MHC-II, F4/80, SiglecF. A representative panel of AMs markers is represented. Histograms represent % of cells iNOS+, Arg-1+, IL-4-r+ and MFI (Mean Fluorescence Intensity) FACS values. Statistical significance is as above.

In addition to their effects on each other’s expression, elafin and IL-6 modify the alveolar macrophage transcriptional profile. Indeed, we show that WT & eTg primary AMs are inherently different : the latter have higher basal levels of the antimicrobial peptides Lcn2 & S100a9 (Fig 1, panels H-I), as well as of the cytokines IL-23 (panel E, trend), IL-10 & IL-22 (panels E-G). Moreover, Ad-IL-6 enhances these differences by up-regulating the expression of these mediators especially Lcn2, S100a9 & IL-10 (panels G-I). Demonstrating specificity, IL-6 and/or Elafin, have no effect on IL-1ß & TNF-a expression (panels J-K).

In addition to the classical AM markers, which did not differ between WT and eTg mice (presence of CD11c, SiglecF & F4/80, Fig1 lower panel, and absence of Ly6C and Ly6G (not shown)), a small, nevertheless significant proportion (10%) expressed the MHC-II molecule. Importantly, elafin and IL-6 expression polarized AMs towards an M2 profile, as demonstrated by an increase in Arg-1+, and IL-4rα+ cell % (Fig. 1 lower panel). Furthermore, elafin and Ad-IL-6 expression in eTg AMs significantly down regulated MHC-II and iNOS MFI levels while up regulating those of IL-4rα. Surprisingly, and in contrast with IL-4rα, Arg-1 MFI was diminished in eTg + Ad-IL6 AMs.

### Adoptive transfer of IL-6- and elafin-genetically modified AMs protect mice against *Pseudomonas aeruginosa*

In order to check the *in situ* behavior of these genetically-modified AMs and their effects on the alveolar environment, 250,000 primary WT- or eTg-AMs, infected with Ad-Null or Ad-IL6, were transferred by oropharyngeal instillation into 7-week old C57Bl/6 male mice and BALs were performed 2 days post transfer to recover alveolar cells (Fig 2A). When compared to controls, all adoptive transfer groups had higher numbers of cells in BALF (2B). Cytospin & MGG staining of the recovered cells revealed a population with macrophagic phenotype without any neutrophilic infiltrate (2D). FACS analysis of these populations showed that they were mostly CD11c+ and possessed comparable surface markers expression among the groups (not shown), except for MHC-II marker that was significantly diminished in eTg-AM + Ad-IL6 transferred group (2C).

**Fig 2.**
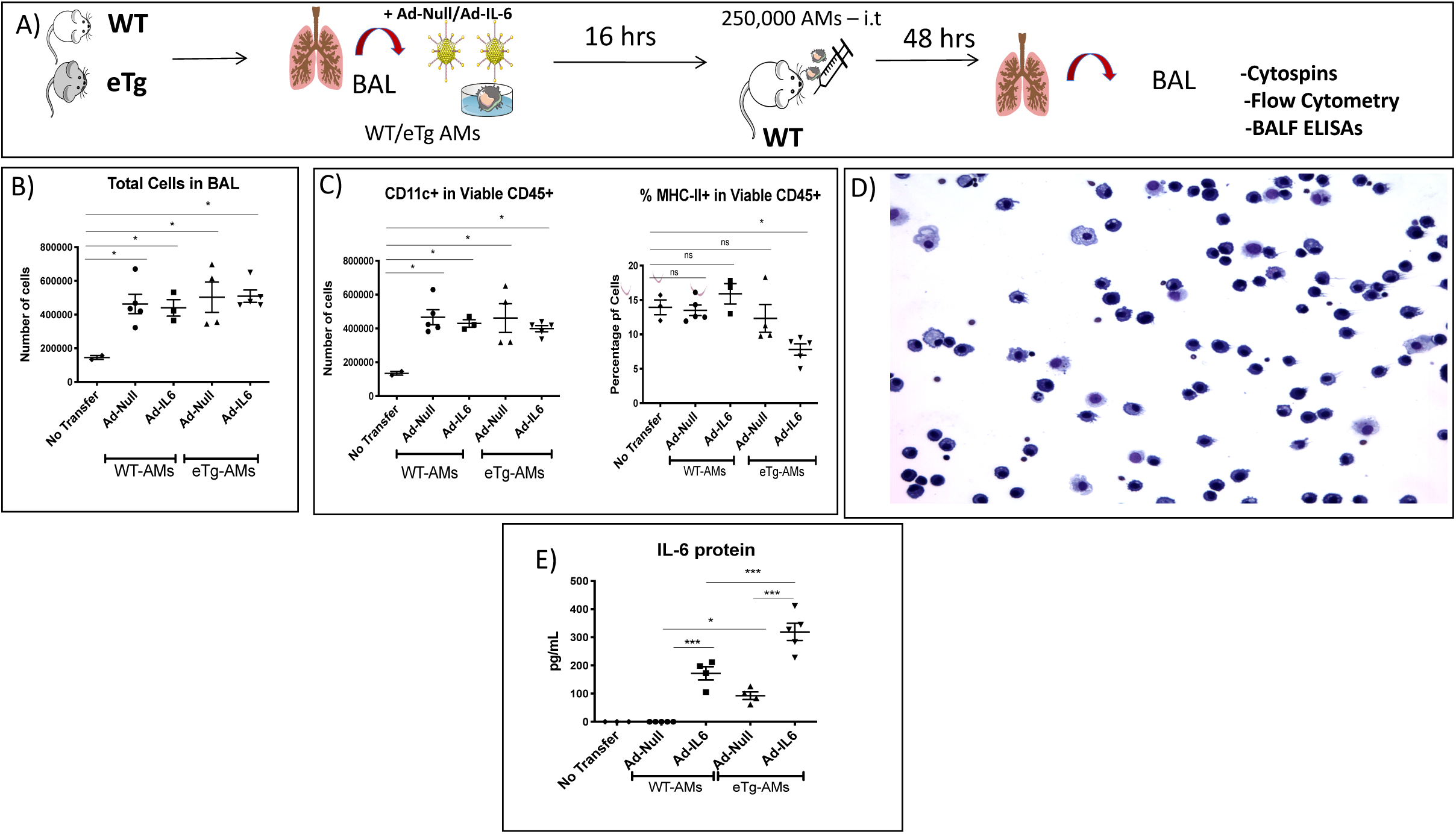
*In situ* characteristics of eTg-AdIL6 AMs adoptively transferred in the lungs of WT C57/Bl6 mice. **A): Experimental procedure:** Primary alveolar macrophages (WT-AM or eTg-AM) recovered by BAL were infected for 6 hrs with adenovirus (MOI 25) Ad-Null (Control-CTRL) or Ad-IL-6 and left overnight. The next day, cells were detached with PBS-EDTA 0,5 mM, washed and the cell pellet was re-suspended in sterile PBS for adoptive transfer by intratracheal route. 250,000 AM was adoptively transferred in each C57Bl/6 mice through the oro-pharyngeal route. 48 hrs post transfer, a BAL was performed, cells counted **(B)**, cytospinned and stained using the MGG Diff Quick staining kit **(D**, a representative BAL is shown). **C: Flow cytometry analysis of BAL cells from adoptively-transferred mice:** CD11c and MHC-II expression in viable CD45+ cells BAL cells is plotted. Protein levels of IL-6 in BALF supernatants is plotted in **E)**. Each point represent an individual mouse and statistical significance is as in Fig 1.

ELISA dosage in BALF (2E) demonstrated higher IL-6 protein levels in the group transferred with eTg-AMs + Ad-null, in accordance with *in vitro* data (Fig 1). Also, expectedly, Ad-IL6 infection significantly increased IL-6 levels in both groups, but the latter were even higher in the eTg-AM group.

Having demonstrated that these adoptively-transferred AMs had retained features of an M2 profile (down-regulation of MHC-II) and were able to produce higher levels of IL-6, we tested these cells both *in vitro* and *in vivo*, in an infectious setting, namely post *PAO1-*infection. *In vitro*, although the differences between WT and eTg AMs were expectedly somewhat attenuated, given the ‘dominant’ huge stimulus conferred by PAO1 infection, some important differences were still noted : IL-23 mRNA and Lcn-2 mRNA levels were still up-regulated by IL-6 (Fig 3, panels A4, A7), and were maximal in eTg AMs, and S100A9 levels were higher in eTg AMs (A8), compared to WT AMs, in the absence of IL-6 stimulation. Interestingly, confirming the ‘M2/regulator’ character of the genetically-modified macrophages, this polarization did not enhance the capacity of primary AMs to clear PAO1 (3B), and even decreased the bactericidal activity of MPI (macrophage cell line) cells over-expressing elafin, alone or in conjunction with IL-6, as well as the production of reactive oxygen species (ROS) production (3C-D). Notably, because MPI murine macrophages do not express elafin (see above), Ad-elafin and Ad-IL-6 constructs were used to over-express these molecules. Importantly, these treatments also biaised MPI responses, in particular through increasing Lcn-2 and S100A9 responses (Fig S1).

**Fig 3.**
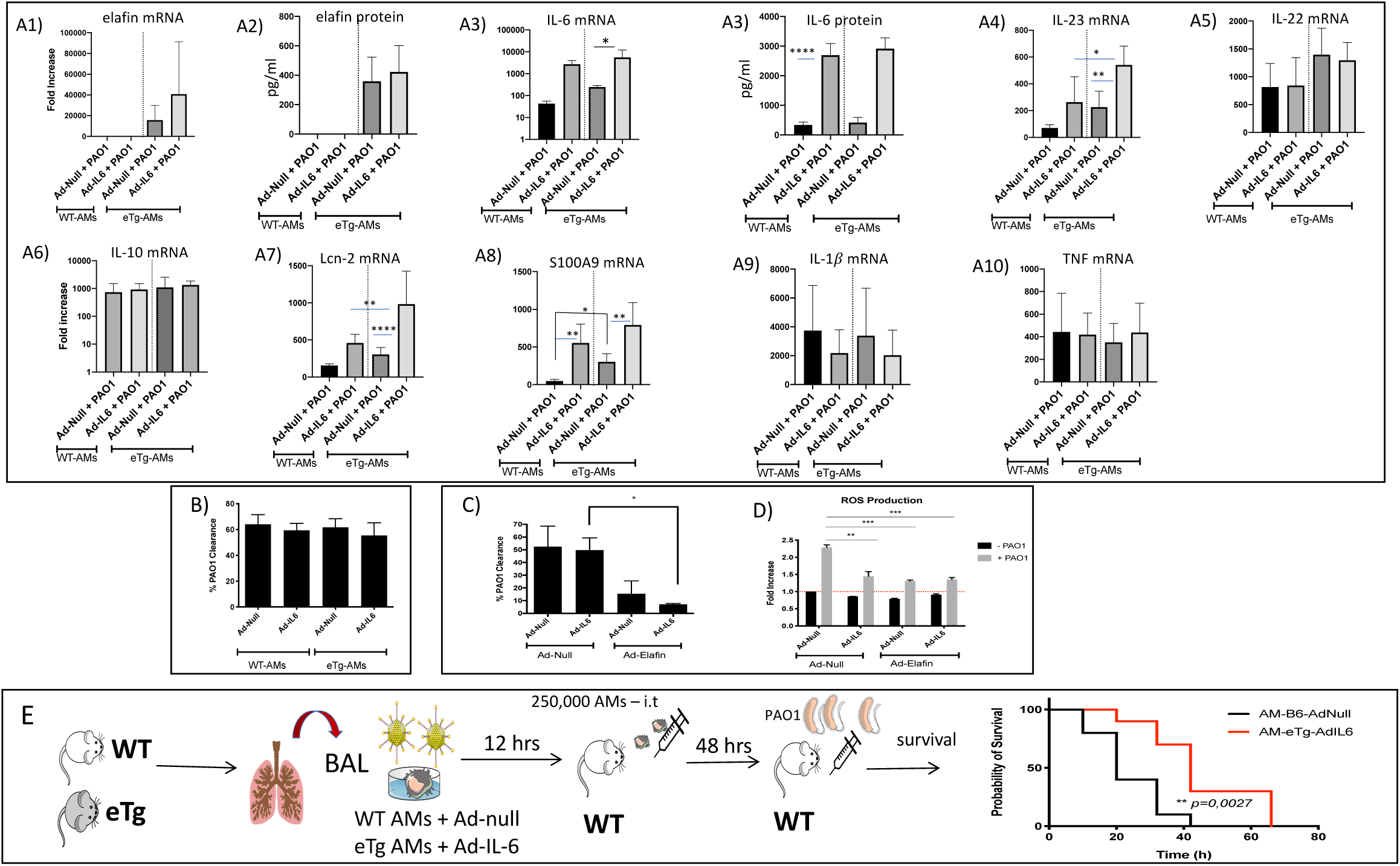
Lung adoptive transfer of eTg-AdIL6 AMs protect mice against *Pseudomonas aeruginosa* (*PAO1*) infection. Panel A: WT or eTg primary alveolar macrophages (WT-AM or eTg-AM) recovered from mice BALs were transfected for 6 hrs with adenovirus (MOI 25) Ad-Null or Ad-IL-6 and left overnight as in Fig 1. The next day, cells were washed and infected with WT PAO1 (MOI 0.5). After 4 hrs, supernatants and cell lysates were treated as in Fig 1 ; Panels B-C: In independent experiments, PAO1 clearance by WT-AM or eTg-AM was measured in WT-AM and eTg-AM (panel B) and in MPI cells, as described in Materials and Methods (panel C) ; Panel D : ROS production in MPI cells transfected with Ad-IL-6/elafin. Statistical significance is as in Fig 1 ; Panel E : After transfer of WT/eTg AMs +/-Ad-IL-6 as in A), C57Bl/6 male mice receivers were infected 48 hours later with 9.10^6 CFU PAO1/mouse. Survival was plotted using Kaplan-Meier curves and statistical tests were performed using the Log-rank (Mantel-Cox) test.

Finally, validating our approach, when comparing the survival of WT C57/Bl6 mice adoptively transferred with either AM-(WT+ Ad-Null) or AM-(eTg + Ad-IL-6) (the two ‘extreme’ conditions) we showed that the latter treatment significantly delayed mortality, after a lethal dose of 10^7 CFU PAO1(Fig 3E).

### Adoptive transfer of IL-6- and elafin-genetically modified BMDMs protect mice against *Pseudomonas aeruginosa*

Because the total number of recoverable AMs is understandingly limited (around 120,000 can be recovered from a naïve mouse), IL-6 and elafin-gene-modified bone marrow derived macrophages (BMDMs) were then isolated from both WT C57/Bl6 and eTg mice, genetically modified as above, and used in most of the rest of the study (Fig 4A).

**Fig 4.**
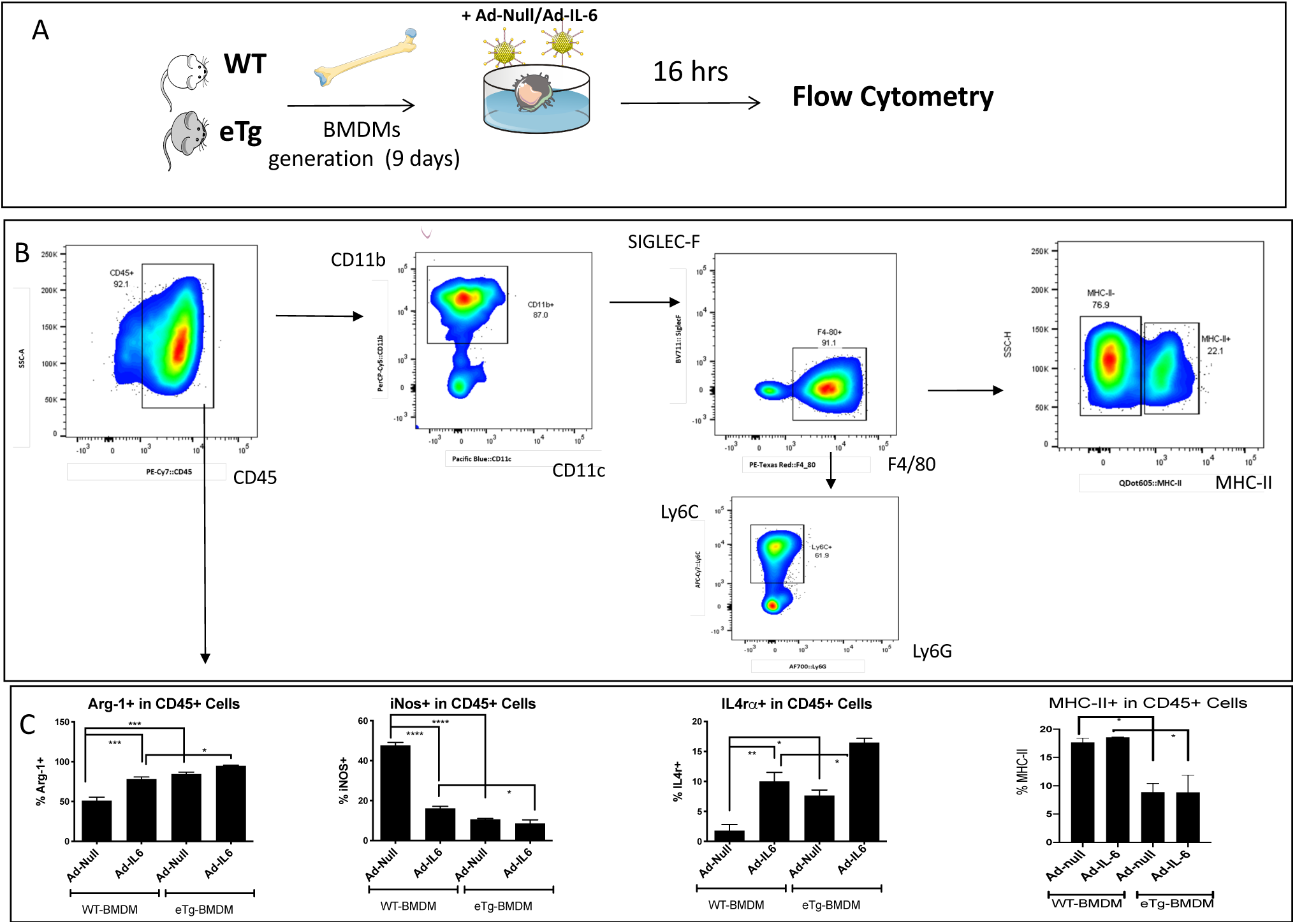
Flow cytometry analysis of BMDMs markers: Elafin and IL-6 expression modify BMDMs expression of Arg-1, iNOS, IL-4r α markers. A) WT and eTg BMDMs were infected for 6 hrs with Ad-Null or Ad-IL-6 (MOI 25). The next day, cells were suspended in PBS-FCS 1% for FACS analysis . B) FACS BMDMs staining procedure, using CD45, CD11b, CD11c, Gr-1 (Ly6c, Ly6G), MHC-II, F4/80, SiglecF markers (a representative Ad-null treatment is depicted. C) FACS study of BMDMs polarization markers : the frequency of iNOS, Arg-1, IL4rα, MHC-II positive cells (among CD45+ viable cells) is depicted. Statistical significance is as in Fig 1

Flow cytometry analysis of WT BMDMs showed that in contrast with primary AMs that are CD11c+ Ly6C-, these cells are CD11b+ Ly6C-/+ (4B). Expectedly, they expressed the surface marker F4/80 that is a hallmark of macrophages, but not the alveolar residency marker Siglec F. A small proportion of BMDMs (around 15%) was MHC-II+, and as found in AMs (Fig 2B), fewer eTg-BMDMs expressed that marker, compared to WT-BMDMs, with or without Ad-IL6 infection, and with (4C). Similarly to what was observed in AMs, IL-6 treatment, in conjunction with the expression of elafin, polarized BMDMs towards an M2 profile, as assessed by increased Arg-1+ and IL-4rα+, and decreased iNOS+ cells frequencies (4C). Interestingly, the elafin-mediated IL-4rα induction was induced at least partly via endogenous IL-6 since an antibody against IL-6 significantly down-regulated it (not shown). Expectedly, *in vitro*, BMDMs produced increased levels of the pro-inflammatory cytokine IL-1b following PAO1 infection (Fig. 5B7-B8), and as shown earlier with primary AMs, basal or PAO1-induced-expression of IL6 was higher in eTg BMDMs than in WT-BMDM (5B2).

**Fig 5.**
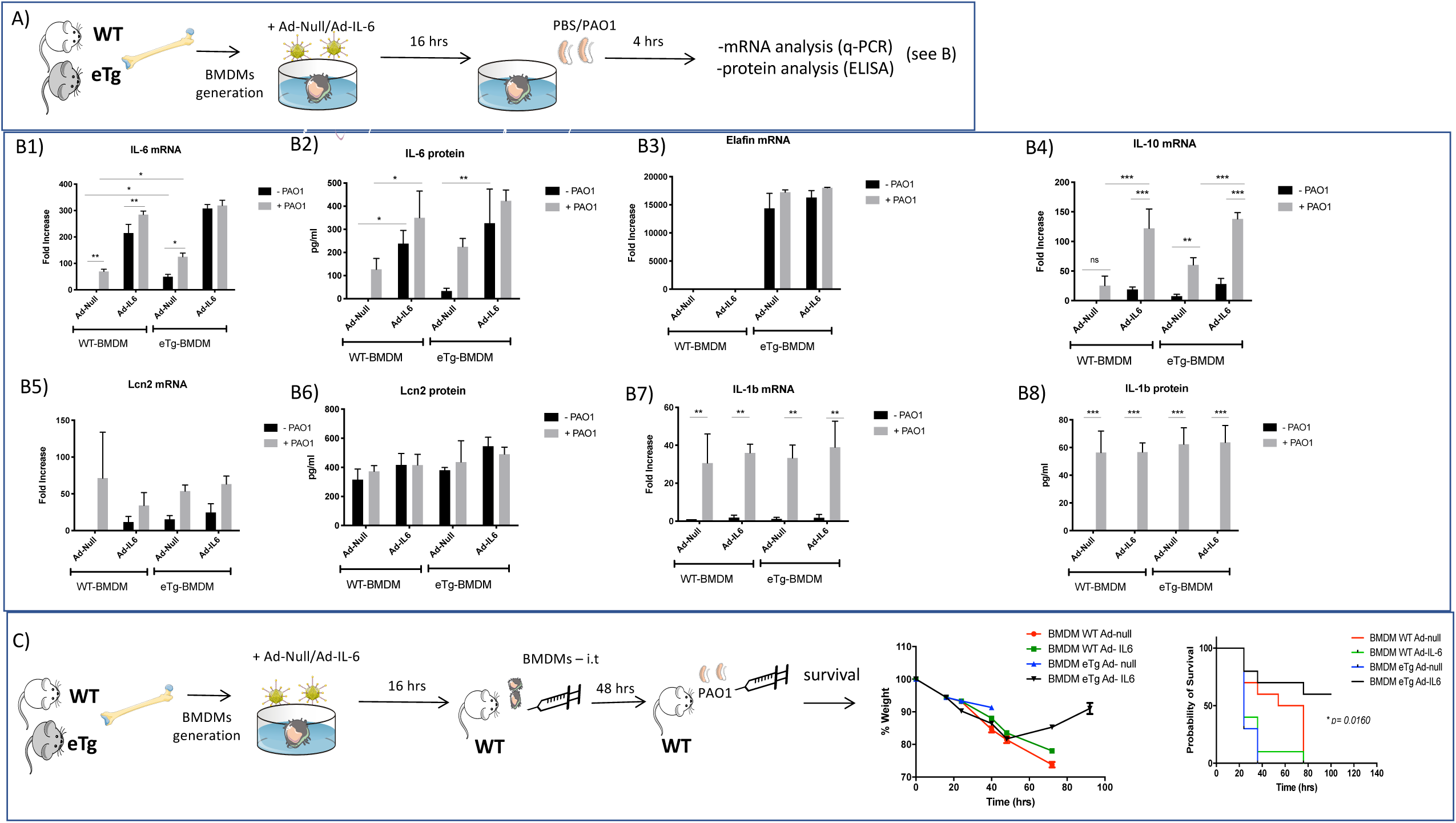
Elafin and IL-6 expression modify IL-10 BMDMs output, and lung adoptive transfer of eTg-AdIL6 BMDMs protect mice against PAO1 infection. A-B) WT and eTg BMDMs were infected for 6 hrs with Ad-Null or Ad-IL-6 (MOI 25) as above, and infected with PAO1 (moi 0.5) for 4 hrs. Then, IL-6, IL-10, IL-1ß, Lcn2 and Elafin mRNA and protein levels were determined by qPCR and ELISA respectively. C) C57Bl/6 WT male mice were transferred with 1,5.10^6 WT/eTg BMDMs pre-infected with Ad-Null or Ad-IL-6, as above. 48 hours after transfer, mice were infected by intranasal route with 9.10^6 CFU PAO1/mouse. Bodyweight loss and survival (Kaplan-Meier curve) were monitored until either animal death or total recovery of the remaining mice. Statistical test was performed using the Log-rank (Mantel-Cox) test.

In order to test the *in situ* behavior of these BMDMs (mock-or genetically-modified), we performed an adoptive transfer (1,5.10^6 / mouse), followed 48hrs later by a lethal PAO1 infection. Our data show that >50% of mice in the eTg-AdIL6-BMDM group survived and were able to recover their initial body weight demonstrating yet again a protective role of elafin and IL-6 in this independent model (5C).

To further investigate the mechanisms explaining the protective effect observed above, we performed a similar experiment of adoptive transfer of WT/eTg BMDMs (genetically modified or not with Ad-IL6), followed 48hrs later by PAO1 infection, and analysed BAL cells and solutes 16 hrs post-infection (Fig 6A).

**Fig 6.**
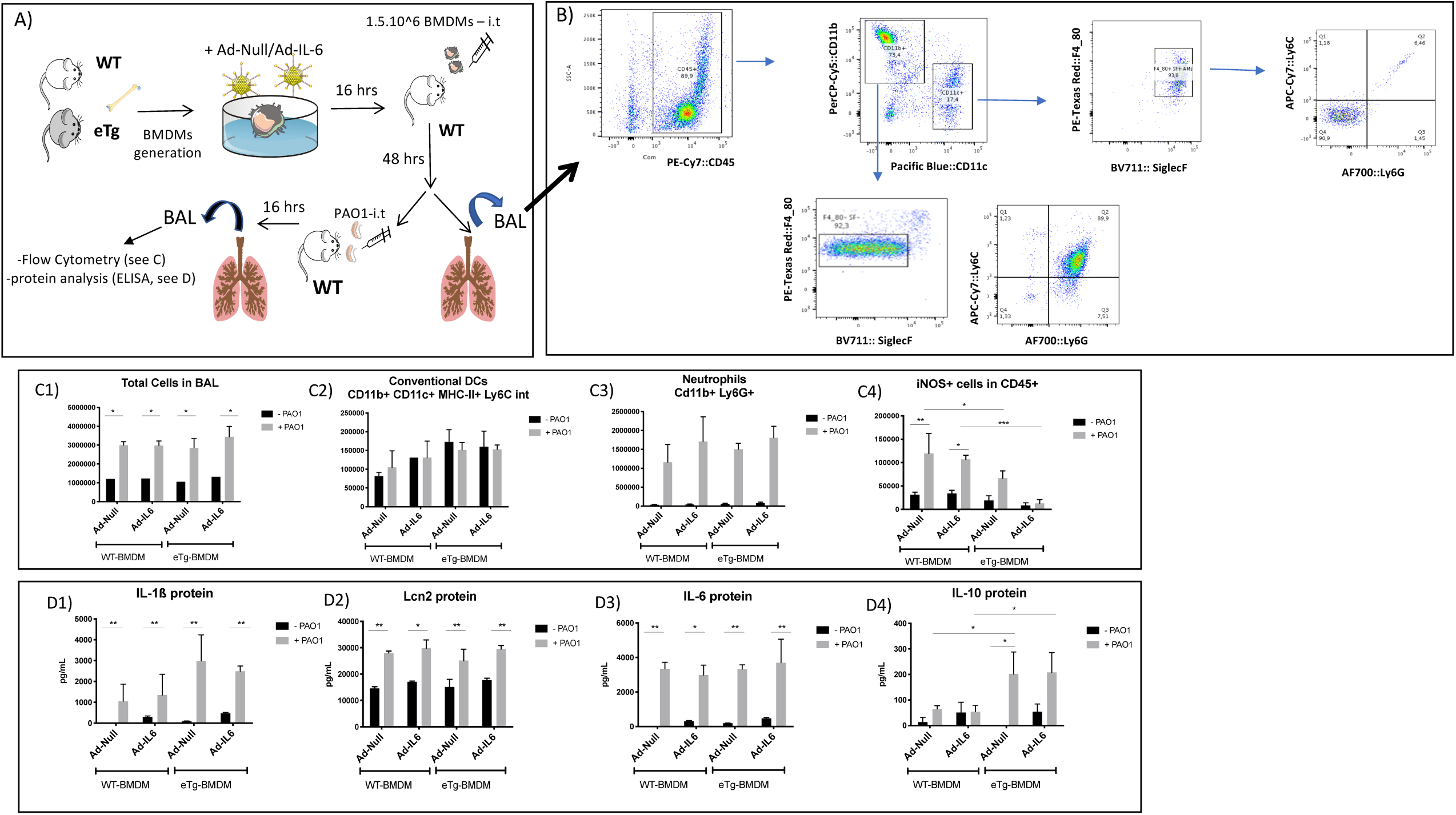
Adoptive transfer of eTg-Ad-IL6 BMDMs induces a regulatory phenotype in the alveolar space upon PAO1 infection. **A)** C57Bl/6 WT male mice were transferred with 1,5.10^6 WT/eTg BMDMs pre-infected with Ad-Null or Ad-IL-6, as above. 48 hours after transfer, a BAL was performed on some mice for FACS analysis of BAL cells **(B**, a representative Ad-null BMDM treatment is depicted). Alternatively, mice were mock- or infected with 6.10^6 PAO1, and BAL was performed 16hrs later for further FACS analysis **(C)** and cytokine protein assessment **(D)**.

FACS analysis of BALs containing unmodified BMDMs at D3 post transfer showed two distinct cell populations (6B). CD11c+ cells (15%) likely correspond to resident AMs since they are Ly6C-, Ly6G-, F4-80+, and SiglecF +, whereas CD11b+ cells most probably include the BMDMs transferred at D0. Interestingly however, the latter became SiglecF+/-intermediate (potentially through the influence of local lung cues ?), and acquired the Ly6G markers becoming double positive for Ly6C and Ly6G, a hallmark of myeloid derived suppressor cells.

Following PAO1 infection, there was, expectedly, an increase in BAL total cell numbers (6C1) in all groups, largely composed of neutrophils infiltrates (6C3) in the alveolar space. No differences were observed in other cell populations in the alveolar space such as dendritic cells (DCs, 6C2). However, iNOS+ cells, which were increased following PAO1 infection, remained low in the eTg-BMDM groups, with or without Ad-IL6 treatment (6C4). Also, BALF IL-1ß, IL-6 and Lcn2 proteins were all induced equally in all groups following PAO1 infection (6D). Interestingly, when compared to WT-BMDM group, the eTg-BMDM group, that had less iNOS+ cells, had high IL-10 levels following PAO1 infection (6D4). This indicates that in addition to low iNOS expression, eTg-BMDM induce the production of high levels of the regulatory / anti-inflammatory cytokine IL-10, suggesting that the high survival rate in this group (Fig. 5) may be the consequence of a dampening of deleterious inflammatory responses.

Because IL-10 is a hallmark of a suppressive environment, we tested the BALF capacity to inhibit T cell proliferation. CFSE-loaded splenocytes were cultured with BALF recovered from the different groups analysed in Fig 6, and activated with CD3/CD28 magnetic beads and left in culture for 72 hours. Flow cytometry analysis of CFSE expression (Fig. 7) shows that BALF recovered from the group transferred with eTg-AdIL6-BMDMs significantly inhibited splenocytes proliferation with or without PAO1 infection, confirming the presence of a suppressive and anti-inflammatory alveolar environment.

**Fig 7.**
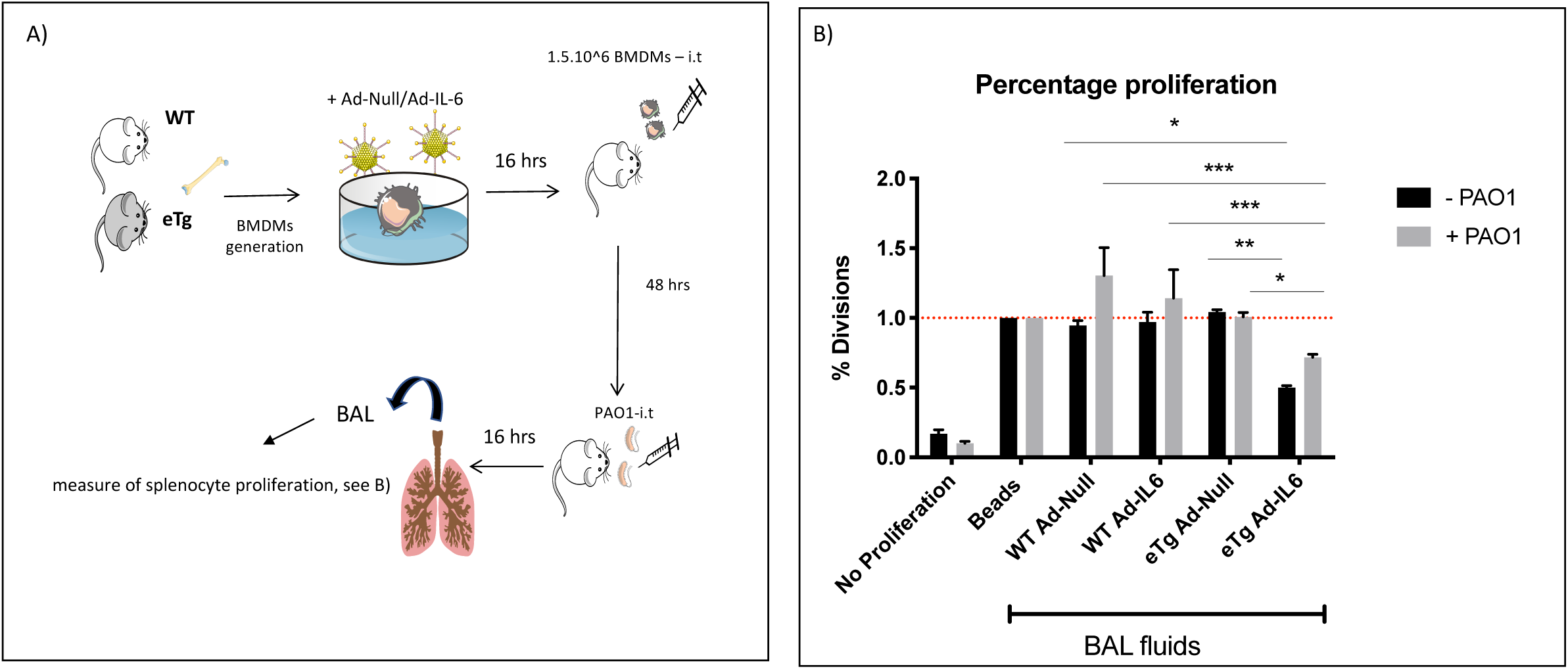
BALF from eTg-Ad-IL6 BMDMs-transferred mice inhibit lymphocytes proliferation *in vitro*. Splenocytes from male WT C57/Bl6 mice were CFSE-labeled and seeded at 10^5 cells/well in a 96 wells plate in DMEM-Glutamax (10% FCS, 1000 U/ml Penicillin, 100 µg/ml Streptomycin) supplemented with 30 U/ml of mouse rIL-2. 2µL of DynaBeads® Mouse T-activator CD3/CD28 were added in each well except for a control well (negative control-no proliferation). 100 µL of PBS “Beads” or BALF recovered from previously transferred and infected mice (Fig 6) were then added into the wells and cells were left in culture for 72 hours. The total number of CFSE+ cells was determined by FACS and the percentage division was normalized to the positive control condition “Beads” given a value of 1 and shown as fold increase.

### BMDM-mediated suppressive phenotype does not rely on T cells or innate-like lymphocytes

Because eTg-IL6-BMDMs are high *in situ* producers of IL-10 (Fig 6) and are able to condition a suppressive alveolar environment (Fig. 7), we wondered whether BMDMs could interact *in situ* with ‘regulatory lymphocytes’. We therefore investigated the effects of this suppressive phenotype during a lethal PAO1 infection in the absence of conventional and/or innate like lymphocytes. To that aim, we performed, as above, adoptive transfer experiments of genetically-modified BMDMs in Rag1-gc double knockout mice. Importantly, following lethal PAO1 infection, we showed that eTg + IL-6 transferred BMDMs still provided protection in that set-up, demonstrating the latter to be independent of lymphocytes (Fig. 8).

**Fig 8.**
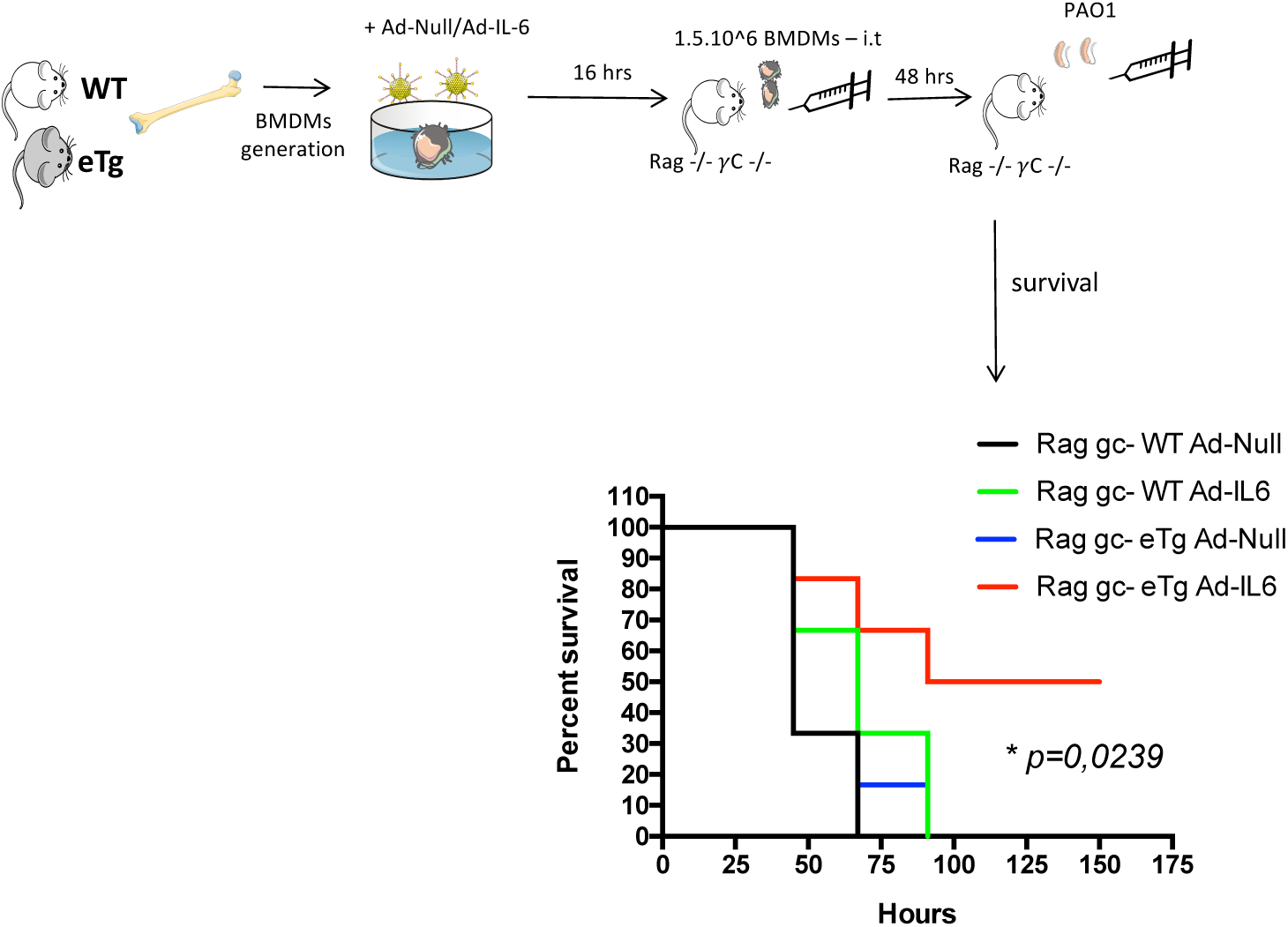
eTg-Ad-IL6 BMDMs-mediated regulatory phenotype does not rely on conventional nor unconventional lymphocytes. Rag gc double KO mice were transferred with 1,5.10^6 WT or eTg BMDMs pre-infected with Ad-Null or Ad-mIL6 (see Figs 6-7). 48 hours post transfer, mice were infected intranasally with 1.10^7 WT PAO1 and survival was monitored until either animal death or total recovery of the remaining mice.

### Adoptive transfer of IL-6- and elafin-genetically modified BMDMs induces a regulatory phenotype in the alveolar space upon PAO1 infection

Because IL-6/elafin BMDM-mediated protection does not occur during the early phase of PAO1-induced response (since pro-inflammatory cytokine levels and cellular infiltrates were identical among the groups, but rather induced a regulatory phenotype (Figs 6-7), we analysed events in a sub-lethal PAO1 infection model at a later time point (instead of stopping the experiment 24 hrs post-infection). To that effect, mice were transferred with either WT-AdNull or eTg-AdIL6 BMDMs (the two ‘extreme’ experimental groups), infected with 5.10^5 CFU of PAO1 and monitored for 5 days, until both groups recovered their original body weight (Fig. 9A-B). In that set-up, differential characterization of BALF cells (9B) showed that eTg-AdIL6-BMDMs group had higher macrophages and lymphocytes numbers and less neutrophils, suggesting a faster recovery process. Elafin mRNA expression in the alveolar cells of this group was increased, indicating that 7 days post BMDM transfer, these cells still remained in the alveolar space (since WT mice do not produce endogenous elafin), despite PAO1-mediated inflammation (9B). Furthermore, ELISA dosage in BALF showed higher levels of IL-6, Lcn-2, Ym-1 and IL-10 in the eTg-IL-6-BMDM-treated groups, suggesting a more efficient lung resolution and repair in this group (9C).

**Fig 9.**
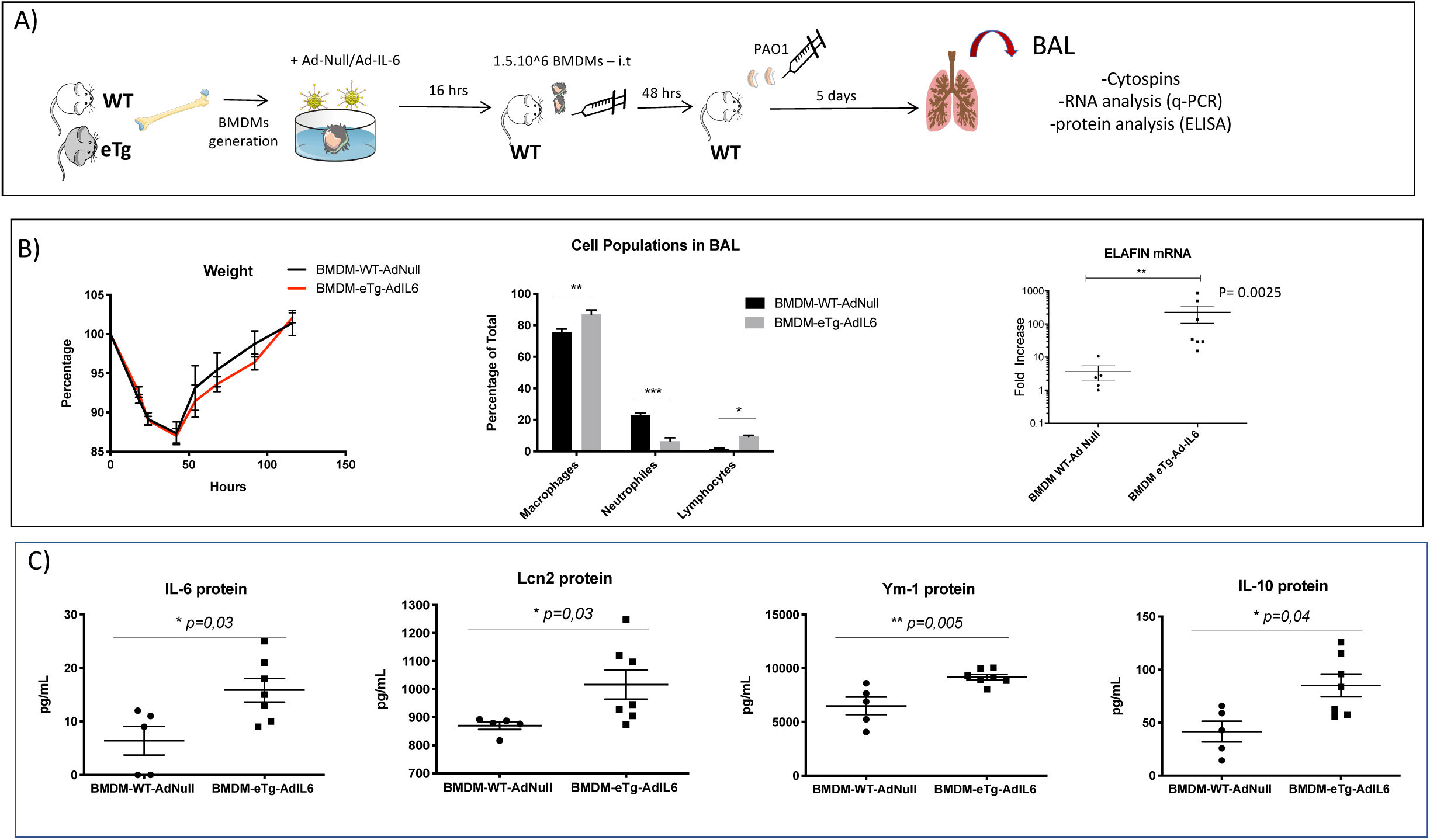
Adoptive transfer of eTg-Ad-IL6 BMDMs enhances a regulatory phenotype in the alveolar space upon PAO1 infection. **A)** Bone marrow derived macrophages (BMDMs) were generated and pre-infected with AdNull or Ad-mIL6 as described in the above Figs. 1,5.10^6 BMDMs were transferred into WT mice and 48 hrs post transfer, mice were infected with a sub lethal dose of PAO1 (5.10^5 CFU/mouse) and euthanized 5 days post infection. **B)** Body weight was monitored daily after PAO1 infection (left panel). At day 5, BAL alveolar cells were counted and differentiated on cytospins (middle panel). BALF pelleted cells were lyzed and Elafin mRNA expression was measured by qPCR. **C)** Recovered BALs were centrifuged and proteins were quantified in BALF by ELISA.

### IL-6 and elafin-modified BMDMs enhance epithelial production of antimicrobial molecules *in vitro* and alveolar epithelial cell proliferation *in vivo*

Because IL-6/elafin BMDM transfer likely transformed the local alveolar environment (see above) independently from lymphocytes, we hypothesized that the nearest targets of AMs / BMDMs would be epithelial cells. Therefore, the interaction between lung epithelial cells and AMs / BMDMs was investigated in a Costar co-culture model *in vitro* where IL-6 and elafin-modified BMDMs and epithelial cells were placed in the upper and lower compartment respectively, and infected (or not) with PAO1 (Fig 10A). Our results show that compared to controls, BMDMs significantly increased mRNA production of the antimicrobial molecule Lcn-2 in both Club cells (DJS-2 cell line, 10B) and alveolar (CMT-2 cell line, 10C) epithelial cells, while S100A9 was up-regulated in Club cells (but completely absent in CMT-2 cells). Furthermore, following PAO1, eTg-BMDMs enhanced Lcn2 expression in CMT but not DJS-2 cells pointing out specificity in Elafin-mediated effects. These Elafin-mediated inductions were up regulated by Ad-IL-6 alone or following PAO1 infection and were somewhat ‘specific’ of antimicrobial molecules since they were not observed with the CCL20 chemokine or IL-6 (the latter result demonstrating that IL-6 does not have an autocrine effect of its own production).

Moreover, in a slightly modified protocol whereby mice were sacrificed 3 days post PAO1 (the onset of tissue repair and epithelial proliferation, 23), we observed that the elafin-IL-6-treated BMDMs induced increased alveolar cell proliferation *in situ*, compared to control-treated BMDMs, upon PAO1 treatment, as assessed by CD45-/Sca-1 positivity (see Fig 10D-E).

**Fig10.**
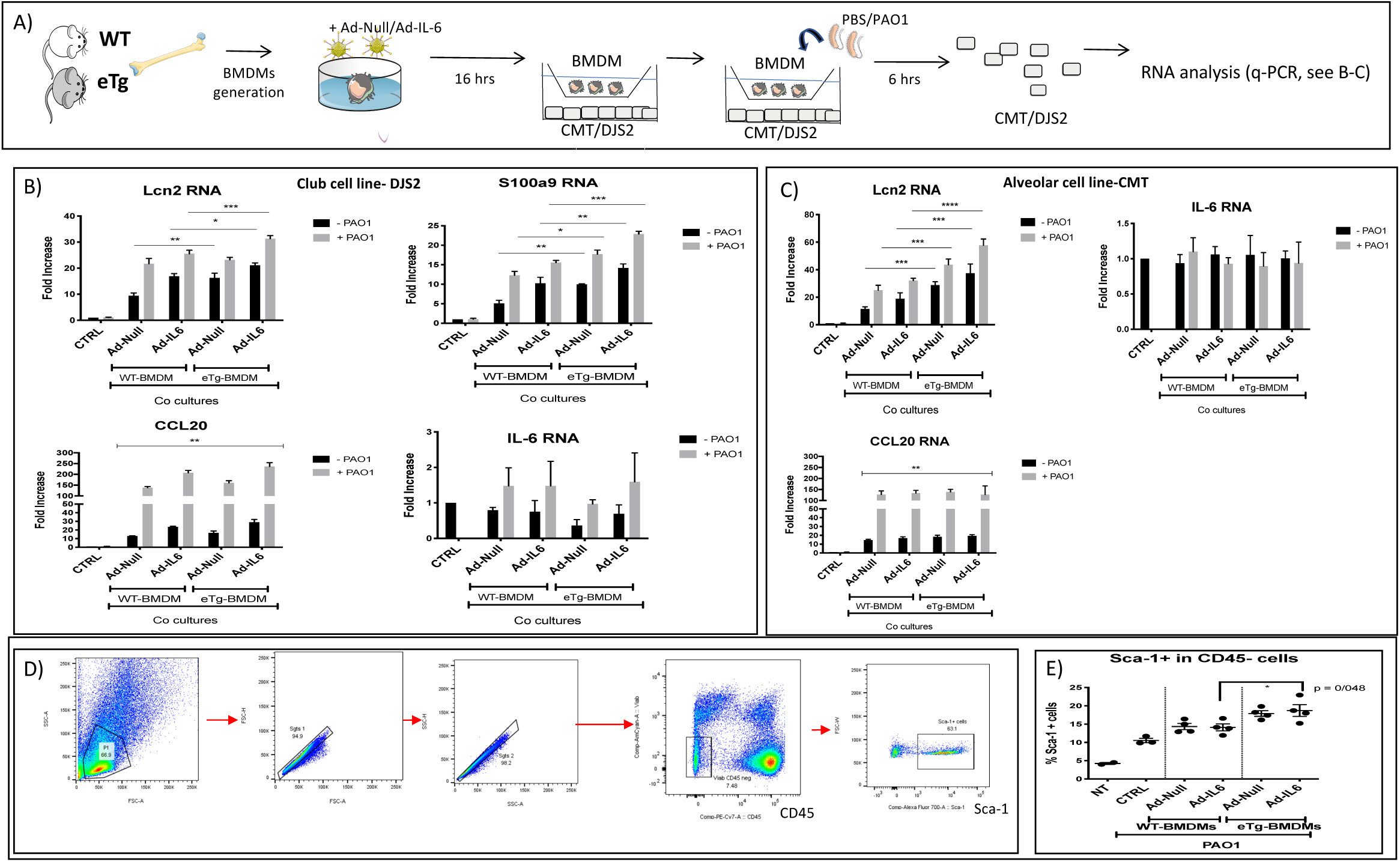
eTg-Ad-IL6 BMDMs enhance epithelial production of antimicrobial peptides *in vitro* and increase epithelial cell proliferation *in vivo*. A) Club cells and alveolar epithelial cells (DJS-2 and CMT-2 cell lines, respectively) were seeded in a 24 well plate (500,000 cells/well). A 12mm diameter insert of 0,4 µM pore size containing 100,000 of either WT or eTg BMDMs (preinfected with Ad-Null or Ad-mIL6) was placed in each insert, except for the non-treated control well (CTRL). After 16hrs, BMDMs were either mock-treated or infected with WT PAO1 (MOI 0,5) for 6 hours. Epithelial cells were then lyzed and antimicrobial peptides / cytokines mRNA expression levels were determined by quantitative RT-PCR. Relative expression normalized to the housekeeping gene 18s of IL-6, CCL20, S100a9 and Lcn2 mRNA in Club **(B)** & alveolar **(C)** epithelial cells ; **(D)** The same protocol depicted in Fig 7 was followed, except that mice were euthanised at day 3, instead of day 5, which better represents the start of the repair process (23). After lung tissue preparation, cells were stained with anti-mouse CD45 and anti-mouse Sca1 antibodies (BioLegend). Flow cytometry analysis: Percentage of Sca-1+ cells within the CD45 negative gate. Viability, CD45+ and Sca-1+ (alveolar proliferation marker) cells are depicted in representative panels; **(E)** The histogram represent % of Sca-1 + in CD45-cells. Statistical significance is as in Fig 1.

### Nuclear elafin transcriptionally up-regulates IL-6 activity

Demonstrating further the direct link between elafin and IL-6 expression in macrophages, we showed that MPI cells, treated with Ad-elafin activated an exogenously added IL-6-luciferase plasmid construct, basally or following PAO1 infection (Fig S2A). Similarly, when MPI cells were treated with exogenously added recombinant elafin proteins (N-elafin, C-elafin, FL-/T2-elafin) all entities (except the N-terminus which does not anti-proteasic activity, see Fig legend) induced strong luciferase activity (Fig S2B), as well as IL-6 and Lcn-2 protein production (Fig S2C). In addition, we demonstrated that elafin recombinant proteins could bind directly to DNA (not shown) and that exogenously added FL- and T2-elafin could be partly recovered in MPI nuclear extracts, suggesting a potential direct stimulatory effect on the IL-6 (and others) promoter (Fig S3).

## Discussion

The lung is constantly bombarded by an arsenal of potentially noxious stimuli such as particles, pollutants and microbes. In that context, maintaining the integrity of the alveolar space is particularly crucial because of the potential frailty of the alveolar-capillary membrane, where 3 cell types interact, at homeostasis, namely the alveolar macrophage (AM) and type I and type II pneumocytes. Indeed, although AMs and epithelial cells pro-inflammatory activities (secretion of interleukin-1, tumor necrosis factor-α, chemokines which recruit and activate inflammatory cells..) are essential in the defense against pathogens [6], [24], [25], it is equally important that the alveolar unit maintain tolerogenic properties at homeostasis, before or after an infectious episode. Indeed, AMs and epithelial cells have been shown to interact closely and provide ‘anti-inflammatory signals’ to the alveolar unit [11], [26-27] and breaking that interaction was shown for example to induce emphysema in a murine model, through alteration of MMP-12 macrophage expression [28].

In that context, we show here that the transfer of a single bolus of syngeneic AMs or BMDMs genetically modified with IL-6 and elafin (eTg) provided protection in a *P.a* lung infection murine model (Fig 3E & Fig 5C). When we mechanistically analysed the modified AMs and BMDMs ‘basally’, prior to transfer, it was evident that the ‘elafin alone’ treatment induced an ‘IL-10/IL-4R’ M2 regulatory signature, when compared to C57/Bl6 WT mice (Figs 1-5). This was associated with decreased MHC-II and ROS macrophage production (a hallmark of ‘M1 macrophages’), and probably relatedly, a ‘neutral’ or even a strong decrease (for MPI cells) in ‘direct’ anti-PAO1 killing (Fig 3B-D).

This signature was reinforced when both elafin and IL-6 were associated, and providing a potential explanation, we showed that these 2 mediators can up-regulate each other expression, and demonstrate for the first time, through independent methods, that elafin can activate the IL-6 promoter (Fig S2). This echoes previous studies indicating that SLPI, a molecule with similar properties, was shown to cross the nuclear membrane thanks to its cationic properties and bind to NF-kB binding sites within the DNA [29]. IL-6 is the archetypical ‘pleiotropic cytokine’, able to promote either pro- or anti-inflammatory responses depending on the context [30-32]. Re-inforcing our data, it has been shown that IL-6 does not on its own bias immune responses, but can, as shown here also, through the up-regulation of the IL4R (Figs 2, 4), enhance the action of IL-4, and hence a M2 phenotype [33-35]. Futhermore, the up-regulation of IL-10 by elafin and IL-6 (Figs 1, 5) is probably also instrumental in up-regulating IL4R, since IL-10 was shown to be a potent inducer of IL-4Ra in a STAT-3 dependent fashion and in return, IL-4 could induce Arg-1 via STAT-6 [36].

Notably, in addition to the ‘typical M2 markers’ mentioned above, elafin expression (alone or with IL-6) was associated or induced the expression of other ‘antimicrobial/regulatory’ molecules such as S100 A9 and Lcn-2. Indeed, although the latter were originally described as antimicrobials [37], it is becoming apparent that their roles, like those of elafin, are wider than originally thought and include anti-inflammatory ones [38-40].

Importantly, the regulatory phenotype described above was maintained *in vivo* post-transfer. Indeed, when the phenotype of un-modified WT BMDMs transfected cells was studied *ex-vivo* at D3 (prior to further *PAO1* infection, Fig 6B), it was apparent that these cells had mostly ‘retained’ a BMDM phenotype (as evidenced by the presence of the CD11b marker), but had however started to ‘convert’ to AMs, as evidenced by the presence of the SiglecF marker (6B), in keeping with the notion that BMDMs can convert *in situ* to AMs [15]. Furthermore, when eTg/IL-6 BMDMs were recovered *ex-vivo* after a further 16hrs PAO1 infection, these cells maintained/promoted a regulatory phenotype in the alveolar space, as evidenced by down-regulation of BMDM iNOS expression (6C), and IL-10 protein accumulation in BALF (6D), which was shown to have *ex-vivo* lymphocyte proliferation properties (Fig 7). This regulatory phenotype was further maintained through day 5, in an independent PAO1-infection experiment where BALs showed increased levels of IL-6, Lcn-2, YM-1 and IL-10 (Fig 9). In addition, we demonstrated, in kinetic experiments involving adoptive transfer of 1,5.10^6 eTg-BMDMs in C57/Bl6 naïve mice (in the absence of IL-6 gene transfer and PAO1 infection), that the number of BAL-recovered cells decreased gradually but remained higher than in non-transferred mice (CTRL) for a period of 2-3 weeks (Fig S4A). During that period, the elafin gene expression remained high (S4C), confirming (since WT mice do not express the elafin gene) that BMDM transferred cells accumulated in the alveolar space and stayed for a long period of time. Furthermore, inflammation was not observed during that time, as evidenced by cytospin analysis (S4B) and measurement of the two pro-inflammatory cytokines MCP-1 and Rantes (S4 E-F). By contrast, IL-6 levels remained sustainable over time (up to 6 weeks, when the experiment was ended), likely demonstrating that eTg BMDMs kept producing elafin and IL-6 for this duration. This may have occurred while adhering to the alveolar epithelium (and thus becoming unrecoverable by BAL, since BAL cells number was ‘back to normal’ by week 3, S4A). Alternatively, it may indicate that despite their turn-over (after 2-3 weeks), eTg BMDMs may have durably modified the alveolar environment to produce high levels of endogenous IL-6, and may have transferred the elafin gene to other local cells, among which probably epithelial cells. Finally, we demonstrated that adaptive or innate lymphocytes were not involved in that protection (Fig 8) and that IL-6 and elafin-modified BMDMs enhanced epithelial production of antimicrobial molecules *in vitro* and alveolar epithelial cell proliferation *in vivo* instead, showing again the importance of lung epithelial cells in the defence against PAO1 (41-42).

In conclusion, although approaches aiming at re-enforcing the protection of the alveolar unit through AM-mediated therapy have been modelled, eg in genetic diseases such as alveolar proteinosis, primary immunodeficiency or alpha-1-Pi deficiency [15], [43-45], less efforts have been geared, as far as we are aware, towards acute infectious situations [16], [46-47]. Our study showing that complementing AMs with a genetic IL-6-elafin combination provide AM with a unique long-lasting new ‘M2-regulatory/antimicrobial’ signature could be of value for treating acute bacterial infections in the lung, i.e in ventilator-associated pneumonia, CF [1], [48] or in lung exacerbations observed in COPD [49].

## Acknowledgements

We wish to thank ‘Vaincre la Mucoviscidose’ for recurrent support, and the Labex ‘Inflamex’ for the PhD studentship award to S. Kheir (grant G11003HH)

## Supplemental material Materials and Methods

### q-PCR Primers

m18S : F : CTTAGAGGGACAAGTGGCG, R : ACGCTGAGCCAGTCAGTGTA ; mTNF : F : AGCCGATGGGTTGTACCTT, R : CAGGGTAATGAGTGGGTTGG ; mLcn2 : F : CCAGTTCGCCATGGTATTTT, R : CCAGTTCGCCATGGTATTTT ; mS100A9 : F : AAAGGCTGTGGGAAGTAATTAAGA, R : GCCATTGAGTAAGCCATTCCC ; mIL-1 ß : F : ATGCCACCTTTTGACAGTGATG, R : GCTCTTGTTGATGTGCTGCT ; mIL-6 : F : GCACCAAGACCATCCAATTC, R : ACCACAGTGAGGAATGTCCA ; mIL-10 : F : AAGGCAGTGGAGCAGGTGAA, R : CCAGCAGACTCAATACACAC ; mIL-22 : F : TTCCAGCAGCCATACATCGTC, R : TCGGAACAGTTTCTCCCCG ; mIL-23 : F : AATCTCTGCATGCTAGCCTGG, R : GATTCATATGTCCCGCTGGTG ; mYM-1 : F : CACGGCACCTCCTAAATTGT, R : CAGGGTAATGAGTGGGTTGG ; h-elafin : F : GGCTCCTGCCCCATTATCT, R : TCTTTCAAGCAGCGGTTAGG

### FACS antibodies

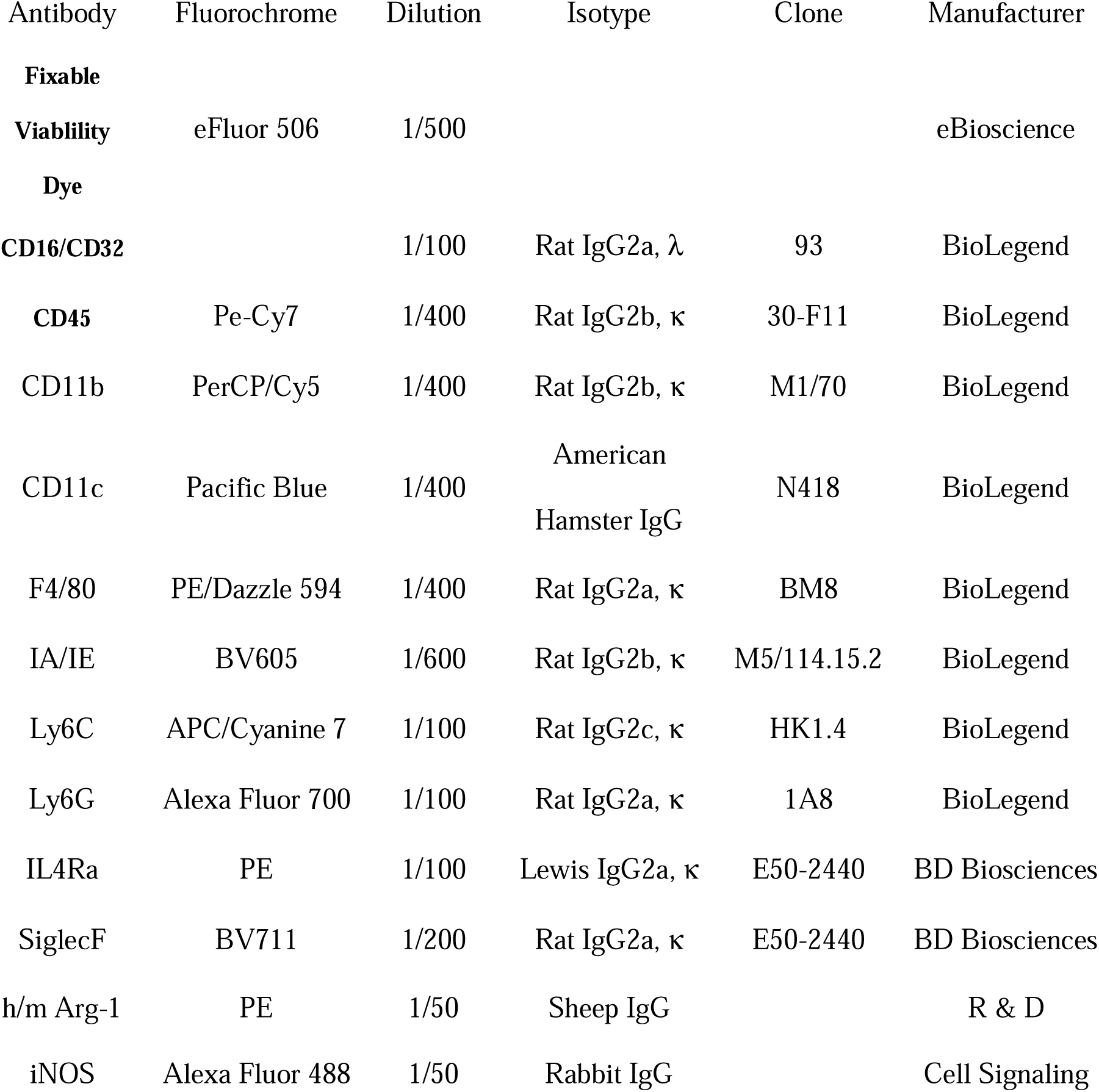

## Results

### IL-6 & elafin over-expression modifies cytokines & antimicrobial peptides production and polarization profile of macrophages of the MPI cell line (Fig S1)

We show, similarly to AMs (Fig 1) that IL-6 and elafin expression was able to polarize AMs. Indeed, Ad-elafin expression increased IL-6, Lcn-2 and S100A9 expression, and Ad-IL-6 over-expression increased elafin expression (Fig S1).

In contrast with primary AMs (Fig 1), macrophages of the MPI cell line have lower expression of antimicrobial molecules (Fig S1 G-H), do not produce IL-10 & IL-22 and respond differently to IL-6 and elafin stimulation : indeed, in that cell line, S100A9 only responds to elafin whereas Lcn2 respond to both IL-6 and elafin, although elafin is still the best ‘stimulus’. Demonstrating specificity, IL-6 and/or elafin, have no effect on IL-1ß & TNF-a expression (Fig. S1, panels E-F).

## Figures legends

**Fig S1: Elafin and IL-6 modify the transcriptional profile of MPI alveolar cells**.

Macrophages (MPI cell line) were infected for 6 hrs with adenovirus (MOI 25) Ad-Null, Ad-IL-6 and/or Ad-Elafin and left overnight. The next day, cells were washed and either mock- or infected with WT PAO1 (MOI 0,5). After a 4-hour infection, cells were lyzed and the cell lysate were used for mRNA quantification by qPCR according to the ΔΔCt method with Relative expression (Fold increase) = 2^^ (-ΔΔCt)^ . The relative expression, normalized to the housekeeping gene 18s, of Elafin and cytokine mRNA, as well as protein levels of IL-6 in supernatants (measured by ELISAs) is presented.

Data are represented as mean +/-SEM. Statistical significance: ANOVA, multiple comparison, Tukey’s test, * : p < 0.05 ; **: p<0.01; *** :p<0.001; ****: p<0.0001).

**Fig S2- Elafin stimulates an exogenous added IL-6-promoter construct and induces IL-6 and Lcn-2 protein production in MPI cells**

**A)** 0.5.10^6^ MPI cells were seeded in 24 well plates containing 0.5 ml of complete RPMI medium (10% FCS, 1% Pen/Strep), and were either mock-treated or infected with either Adnull or Ad-Elafin at a MOI of 25, as described in the Main M&M. After an o/n incubation, cells were further transfected with 1ug of plasmid DNA coding for the IL-6 promoter 5’ of the luciferase gene (pmIL-6 FL #61286-Addgene), diluted into 0.25ml of OptiMEM medium, to which 0.25 ml of OptiMEM 3% lipofectamine 2000 was added. After 6hrs, cells were washed and replenished with complete RPMI overnight. 16hrs later, cells were washed again and were either mock- or infected with PAO1 (moi 1) for 5hrs in serum-free RPMI. At the end of the experiment, cells were lysed for assessment of RLI: luciferase activity (Promega kit). UN : untransfected cells, i.e no Ad-, nor plasmid pmIL-6 FL were added. Cells transfected with pmIL-6 FL alone exhibited the same RLI as cells transfected with both Adnull and pmIL-6 FL ; Ad-null : cells transfected with both Ad-null and pmIL-6 FL ; Ad-Elafin : cells transfected with both Ad-null and pmIL-6 FL

**B-C)** 0.5.10^6^ MPI cells were seeded and transfected with the IL-6 promoter-luciferase construct as in **A)**. Cells were then incubated during 5hrs in serum-free medium with increasing concentrations of 0.4 uM, 2 uM, 4uM of FL (full-length)-elafin (a 95 aa molecule chemically synthesized, with a net cationic charge of +7 and active as an anti-protease, (48) or with the same concentrations of trappin-2 (T2-elafin (same as FL-elafin, but obtained commercially (R&D)). Alternatively, cells were incubated with 0.8uM, 4 uM, 8uM of C-terminus elafin (aa 38-95 of FL-elafin, with a net cationic charge of +3 and active as an anti-protease) or with the same concentrations of N-elafin (aa 1–50 of FL-elafin, with a net cationic charge of +5 and inactive as an anti-protease). At the end of the experiment, supernatants were recovered for measurement of IL-6 and Lcn-2, see

**C)** (R&D ELISA commercial kits) and cells lysed for assessment of luciferase activity, see **B)** (Promega kit).

Data are represented as mean +/-SEM. Statistical significance: ANOVA, multiple comparison, Tukey’s test, * : p < 0.05 ; **: p<0.01; *** :p<0.001; ****: p<0.0001).

**Fig S3 Exogenously-added elafin penetrates MPI cells and a proportion localizes in the nucleus**.

0.5.10^6 MPI cells were seeded in 24 well plates containing 0.5 ml of complete RPMI medium (10% FCS, 1% Pen/Strep) and left overnight. The next day, cells were PBS-washed and incubated with increasing concentrations (0.4uM, 2uM, 4uM) of FL-elafin or T2-elafin (see Fig S3) in serum-free RPMI medium for 1 hour. At the end of the experiment, cells were lysed and subcellular protein fractionation was performed using the NE-PER Nuclear and Cytoplasmic Extraction reagents (Thermo Scientific). Cytoplasmic **(A)** and nuclear **(B)** Elafin concentrations were determined by ELISA). Data are represented as mean +/-SEM

**Fig S4 BMDMs kinetics in the alveolar space following intra-tracheal adoptive transfer**

Elafin transgenic (eTg) BMDMs were generated as described in the Materials and Methods and transferred intra-tracheally into WT C57/Bl6 mice (1.5.10^6^ /mice) as explained above. Mice were then euthanized at different time points post-transfer (Day 0, weeks 1, 2, 3, 6) and alveolar cells were recovered by BAL. **A)** Number of total cells recovered in BALF ; **B)** Cells recovered in BALF at week1 were stained using the MGG Diff Quick staining kit ; **C-F)** Elafin, IL-6, RANTES and MCP-1 mRNA levels were determined by q-PCR in recovered cells. Data are represented as mean +/-SEM, and each point represent an individual mouse. Statistical significance: **A)** ANOVA, multiple comparison, Tukey’s post-hoc test test, ****: p<0.0001) ; **C-F)** : Kruskal-Wallis, multiple comparison, Dunn’s post-hoc test, * : p < 0.05

