## Supplementary figures and images for "Use of IL-6-elafin genetically modified regulatory macrophages as an immunotherapeutic against acute bacterial infection in the lung"

### Figs S1-S4

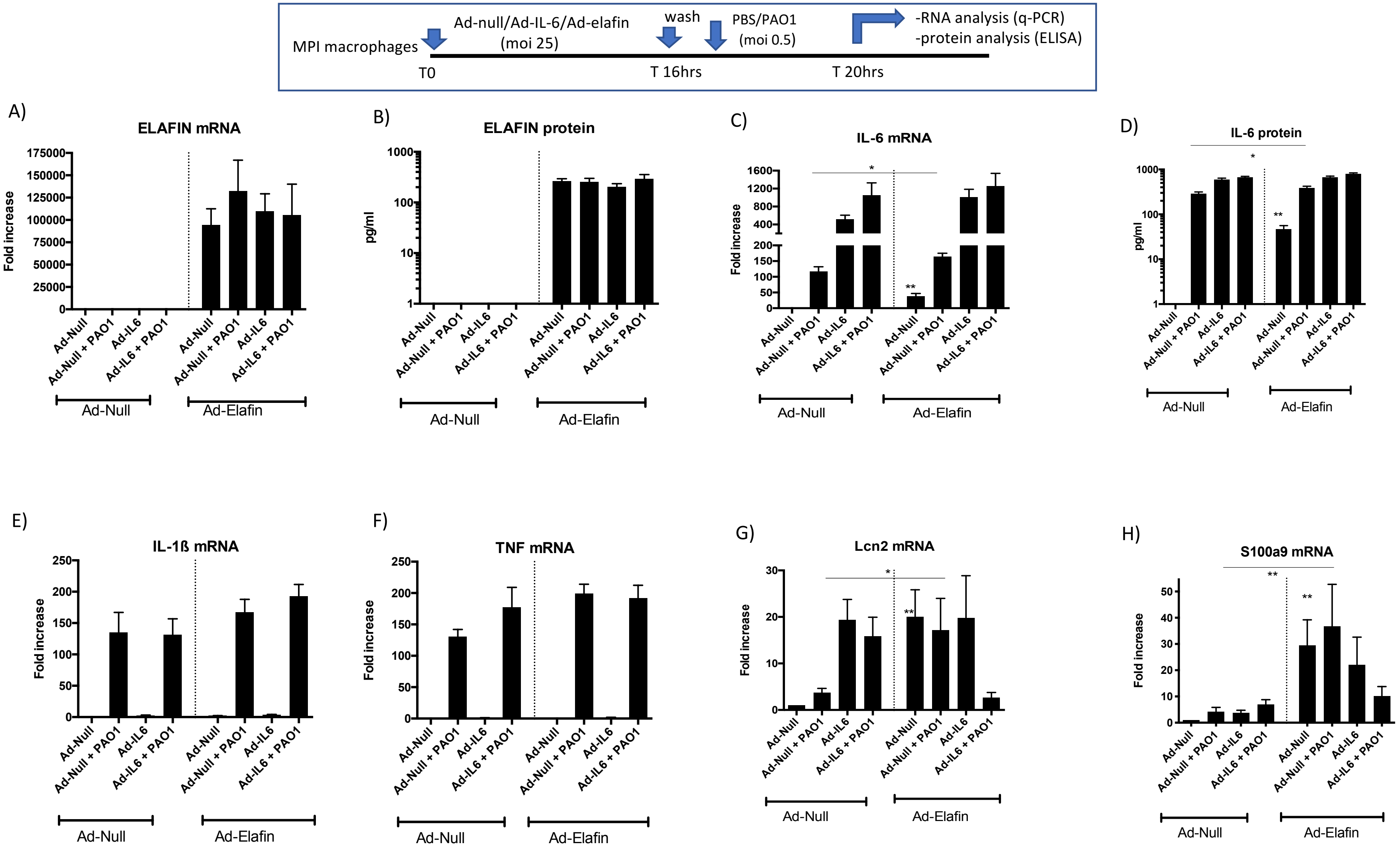

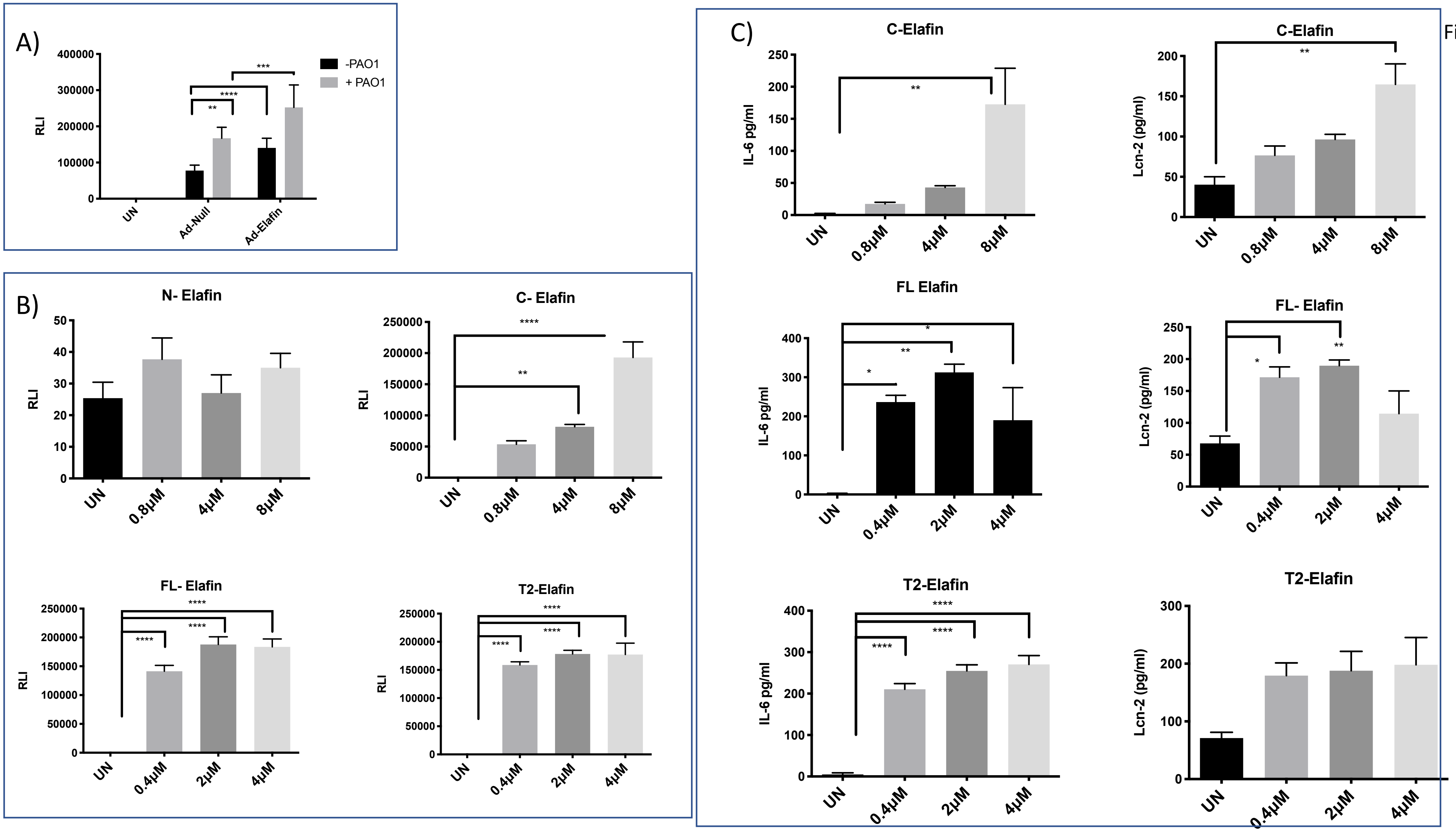

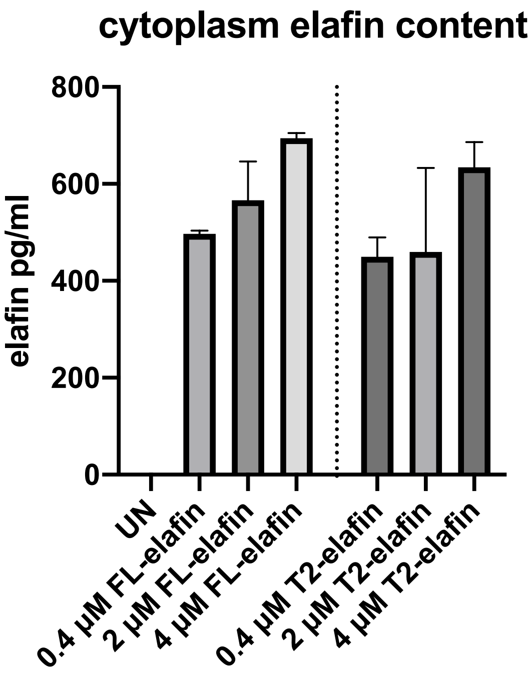

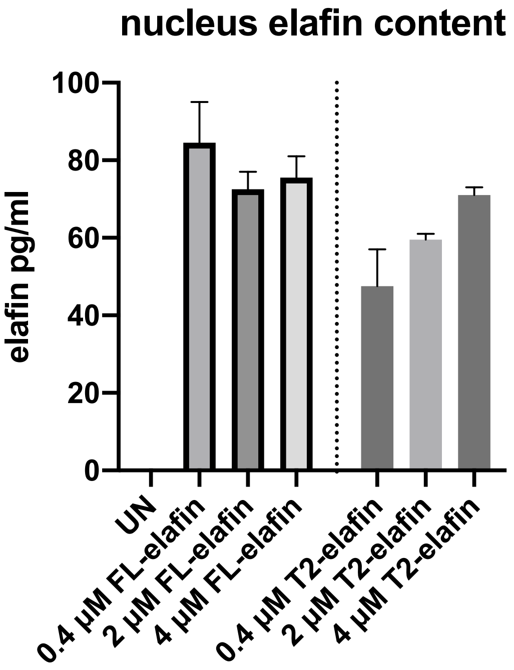

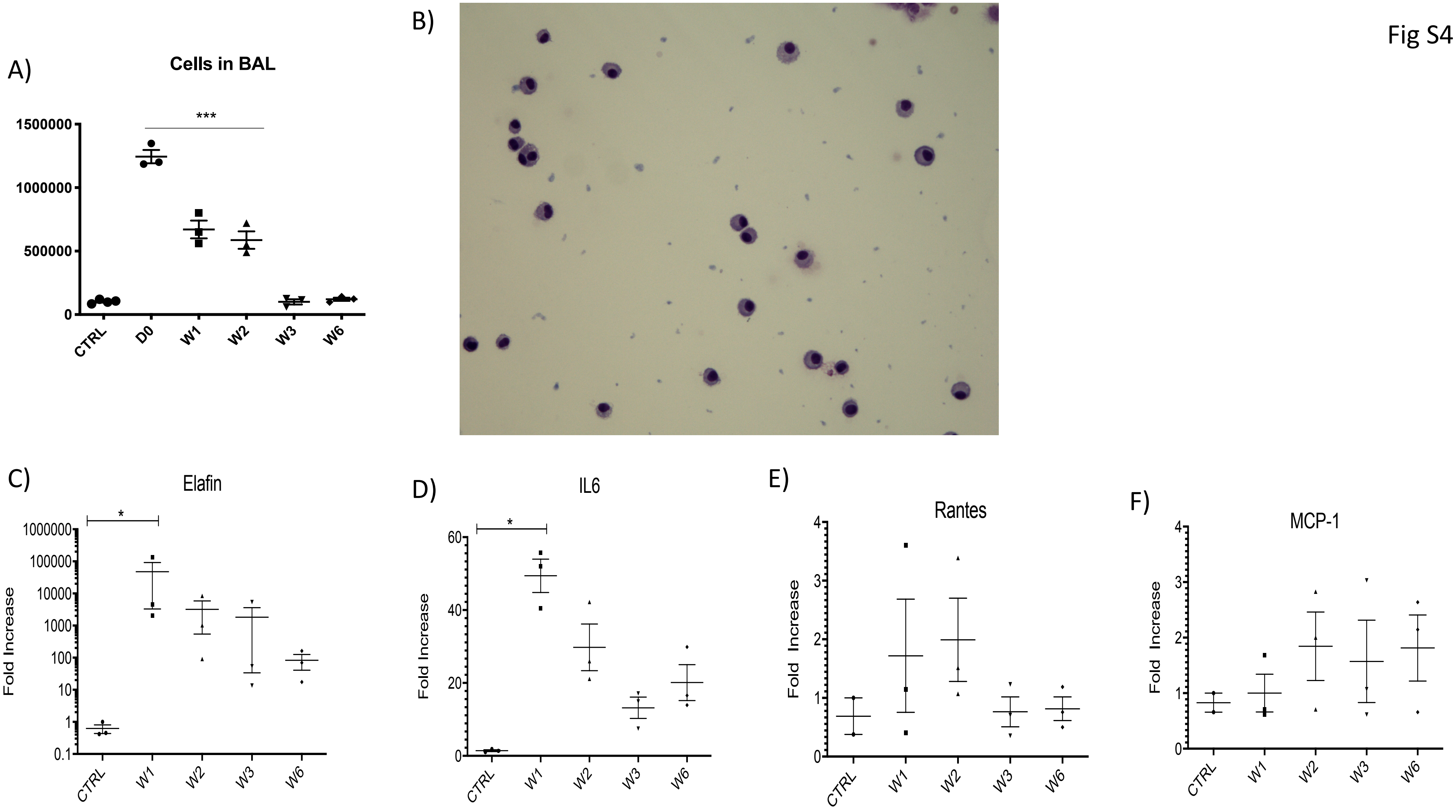
